# Carbonic anhydrase inhibitors prevent presymptomatic capillary flow disturbances in a model of cerebral amyloidosis

**DOI:** 10.1101/2024.08.22.609091

**Authors:** Eugenio Gutiérrez-Jiménez, Peter Mondrup Rasmussen, Irene Klærke Mikkelsen, Sreekanth Kura, Signe K. Fruekilde, Brian Hansen, Luca Bordoni, Jasper Carlsen, Johan Palmfeldt, David A. Boas, Sava Sakadžić, Sergei Vinogradov, Mirna El Khatib, Jaime Ramos-Cejudo, Boris Wied, Desiree Leduc-Galindo, Elisa Canepa, Adam C. Mar, Begona Gamallo-Lana, Silvia Fossati, Leif Østergaard

## Abstract

**INTRODUCTION:** Disturbances in microvascular flow dynamics are hypothesized to precede the symptomatic phase of Alzheimer’s disease (AD). However, evidence in presymptomatic AD remains elusive, underscoring the need for therapies targeting these early vascular changes.

**METHODS:** We employed a multimodal approach, combining in vivo optical imaging, molecular techniques, and ex vivo MRI, to investigate early capillary dysfunction in Tg-SwDI mice without memory impairment. We also assessed the efficacy of carbonic anhydrase inhibitors (CAIs) in preventing capillary flow disturbances.

**RESULTS:** Our study revealed capillary flow disturbances associated with alterations in capillary morphology, adhesion molecule expression, and Amyloid-β (Aβ) load in 9–10-month-old Tg-SwDI mice without memory impairment. CAI treatment ameliorated these capillary flow disturbances, enhanced oxygen availability, and reduced Aβ load.

**DISCUSSION:** These findings underscore the importance of capillary flow disturbances as early biomarkers in presymptomatic AD and highlight the potential of CAIs for preserving vascular integrity in the early stages of AD.

## 1. Background

Alzheimer’s disease (AD) is the leading cause of dementia globally, affecting 46.8 million in 2015^1^. In Europe, AD prevalence is predicted to double by 2050^2^, underscoring the urgent need for effective disease-modifying interventions. However, the precise biological causes of AD remain unclear. AD is understood as a biological and clinical continuum that develops gradually over years, marked by pathological changes such as Amyloid β (Aβ) plaques, cerebral Aβ angiopathy (CAA), hyperphosphorylated tau protein aggregation, and neurodegeneration^3^.

Supporting the continuum model of AD, human biomarker studies suggest that Aβ accumulation begins at least 15 years before clinical symptoms^4^. While therapeutic strategies targeting Aβ effectively reduce Aβ load, they have shown limited success in controlling disease progression and fail to reverse memory loss^5,6^. This highlights the need to identify and target early disease mechanisms manifesting long before symptoms arise.

AD pathology and cerebrovascular disease (e.g., CAA) frequently coexist in the elderly, with recent studies highlighting a potential synergistic relationship^7^. Vascular contributions to AD are supported by changes in brain vasculature detected up to 10 years before symptomatic AD and possibly before Aβ deposits^8^. AD also shares many risk factors with cardiovascular diseases^9,10^. Evidence from animal models shows vascular changes involving disturbed neurovascular coupling^11^, blood-brain barrier (BBB) dysfunction^12^, vascular oxidative stress, and inflammatory damage^13^, ultimately reducing cerebral blood flow and limiting oxygen and nutrient access to brain tissue. Regulating local cerebral blood flow (CBF) is essential for local oxygen delivery. Capillary transit time and blood flow distribution across the capillary bed significantly influence oxygen extraction^14^. Disturbances in capillary flow dynamics including transient blockage by leukocytes or platelets, known as capillary stalls^15^, and capillary constrictions due to Aβ plaques^16^ have been observed in animal models of AD. These disturbances may increase capillary resistance, disrupt flow distribution, and reduce oxygen extraction efficiency, termed capillary dysfunction^17^.

Recent observations highlight the importance of early capillary flow disturbances in AD pathology^17^. Elevated capillary transit-time heterogeneity (CTH) is observed in AD patients and individuals with mild cognitive impairment^18^ and at preclinical stages^19^. Similarly, a murine AD model exhibits increased CTH in symptomatic stages^11^. However, capillary flow disturbances have not yet been observed during presymptomatic stages of AD.

Therapeutic means to prevent the deterioration of capillary function may be crucial to prevent AD. Recent studies highlight carbonic anhydrase inhibitors (CAIs) as potential AD treatments^20,21^. CAIs are clinically used in treating glaucoma^22^ and mountain sickness^23^. CAIs inhibit the reversible hydration of carbon dioxide (CO_2_) to bicarbonate and an hydrogen proton, a process playing a critical role in maintaining pH homeostasis^24^. Elevated CO_2_ levels enhance CBF through vasodilation^21^, while the resulting metabolic acidosis right-shifts the haemoglobin-oxygen dissociation^25^. Both these actions therefore improve tissue oxygenation. Prior studies also show that CAIs reduce Aβ oligomer neurotoxicity by preventing mitochondrial dysfunction and pro-apoptotic mechanisms^20,21,26^. We recently showed that in 15-month-old Tg-SwDI mice, long-term CAI treatment with Methazolamide (MTZ) or Acetazolamide (ATZ) prevented cognitive decline, reduced vascular and total Aβ load, improved glial clearance, and limited inflammation^27^. However, no studies have so far examined the effect of CAIs on brain microcirculation in vivo in mouse AD models.

This study aims to determine whether early capillary flow disturbances exist in presymptomatic Tg-SwDI mice, a model of cerebral amyloidosis, and whether these alterations lead to capillary dysfunction (increased CTH), and impaired oxygen uptake. Additionally, it examines whether early CAI treatment mitigates capillary flow disturbances in presymptomatic Tg-SwDI mice, thereby improving oxygen availability, Aβ load, or brain structure.

Using a multimodal approach, we integrated in vivo optical imaging, molecular biology, and ex vivo MRI to investigate early capillary dysfunction in Tg-SwDI mice and the impact of CAI treatment. The Tg-SwDI model represents the AD continuum, with Aβ accumulation starting at 4 months without significant symptoms until 12 months^28^. We demonstrate capillary hemodynamic disturbances in 9-10-months-old presymptomatic Tg-SwDI mice. Early treatment, particularly with MTZ, significantly prevent these capillary flow disturbances, enhances oxygen transport, and prevent Aβ load. Our findings suggest that CAIs, particularly MTZ, may prevent vascular impairment and slow down AD pathology during presymptomatic stages.

## 2. Methods

### 2.1 Experimental model

The Danish Ministry of Justice and Animal Protection Committees approved all animal procedures and treatment, with the Danish Animal Experiments Inspectorate permit 2017- 15-0201-01241. We used adult homozygous mice from the Tg-SwDI line (C57BL/6- Tg(Thy1-APPSwDutIowa)BWevn/Mmjax; MMRRC Stock No: 34843-JAX, The Jackson Laboratory). A group of homozygous breeders was acquired from Jackson Laboratory and bred at the animal facility at the Biomedicine Department, Aarhus University). Animals were of both sexes. A group of control wild-type mice (C57BL/6J) were acquired from Jackson Laboratory, matching sex and age. After weaning from the breeder, mice were transported to the stable facility at the Center of Functionally Integrative Neuroscience (CFIN, Aarhus University). Mice were housed in group cages (between 3 - 5 mice/cage) with *ad libitum* access to water and a standard diet/treatment diet. Mice maintained at 12h:12h light-dark cycle at constant temperature and humidity (21 °C ± 2 and 45% ± 5 relative humidity). Behavioral studies were performed on mice with identical group characteristics, at NYU School of Medicine Rodent Behaviour Laboratory and adhered to the guidelines of the NYU School of Medicine’s Institutional Animal Care and Use Committee as in previous work^27^.

To ensure sufficient statistical power, we conducted a power analysis based on previous work by Park et al., which demonstrated cerebrovascular disturbances in the Tg-SwDI^29^. Two reference age groups, 3 months and 18 months, were used to estimate the cerebrovascular response metrics for the 9–10-month cohort. The 9-month sample data were calculated as the midpoint between the 3-month data (WT: mean = 23 ± 1 SEM, Tg- SwDI: mean = 16.5 ± 1 SEM) and the 18-month data (WT: mean = 13 ± 1.5 SEM, Tg- SwDI: mean = 9 ± 1.5 SEM) for the CBF response to whisker stimulation, as described in Figure 1A of Park et al. The initial power analysis indicated a required sample size of 6 mice per group to achieve 80% power with a significance level of 0.05. However, as our experimental design involves awake mice, where we anticipate a 60% success rate in completing experiments, we adjusted the sample size to account for potential losses. After this adjustment, the required sample size increased to 10 animals per sex per group.

**Figure 1.**
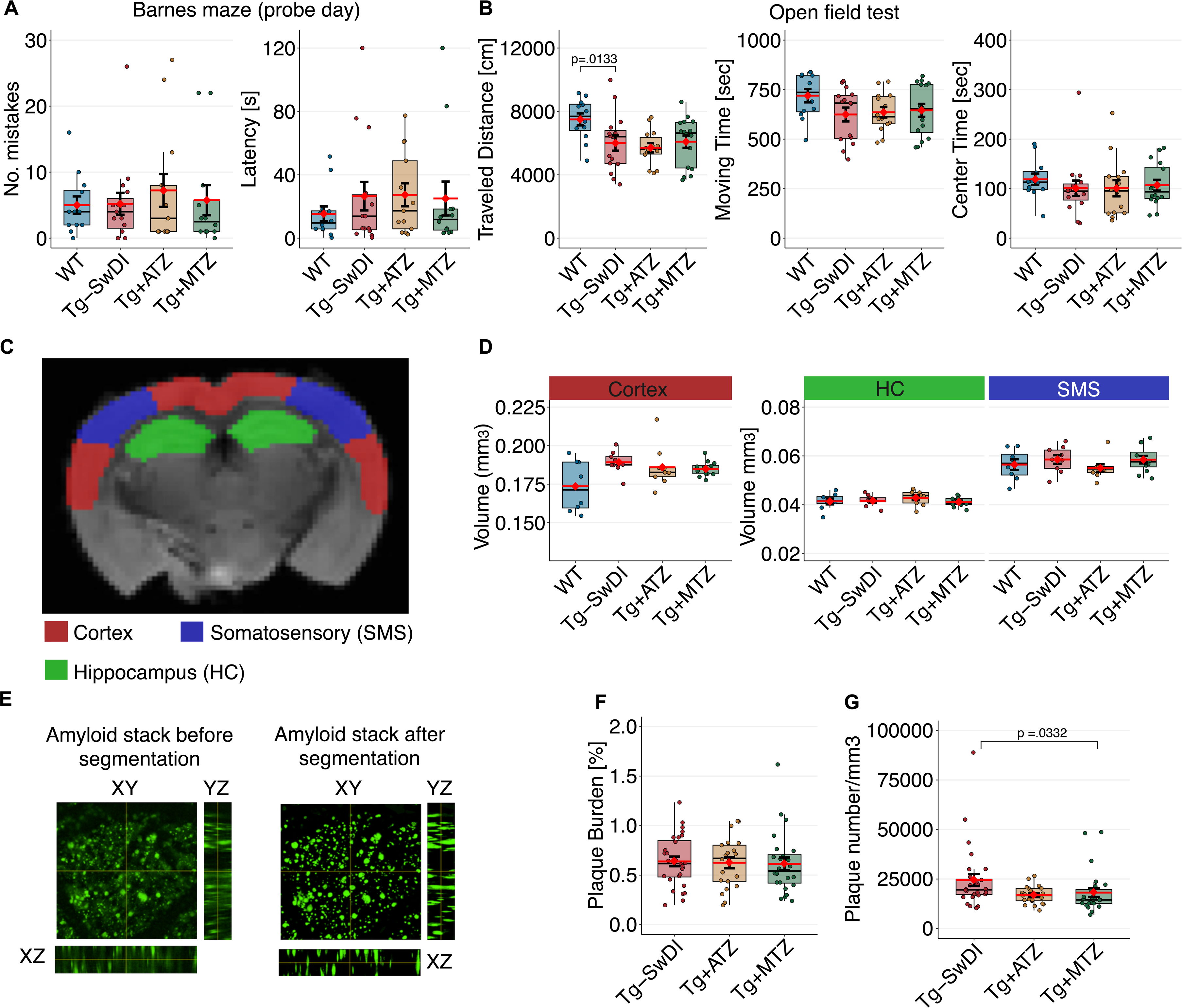
Early amyloidosis and cortical hypertrophy in cognitively normal TgSwDI mice. **A).** Analysis of Barnes maze in WT and Tg-SwDI mice with and without treatment. WT n=12 (6 female (F), 6 male (M)); Tg-SwDI n=15 (5 F, 10 M); Tg+ATZ n=13 (5 F, 8 M); Tg+MTZ n=12 (6 F, 6 M). Plots represent the number of mistakes (left), and latency (right). **B)** Analysis of open field to assess locomotor activity. WT n=12 (6 F, 6 M); Tg-SwDI, n=15 (5 F, 10 M); Tg+ATZ, n=14 (6 F, 8 M); Tg+MTZ, n=16 (8 F, 8 M). Plots represent traveled distance (left), moving time (middle), and time spent on field’s center (right). Traveled distance across groups (ANOVA: F[3, 53] = .337, p = .798). WT vs Tg-SwDI pairwise comparisons using EMMs, diff = –1493 ± 583 cm, t(53) = −2.561, d = .906. **C)** Regional atrophy was assessed estimating regional tissue volume using ex-vivo MRI. Segmentation into cortex, hippocampus (HC), somatosensory area (SMS), and white matter (WM) was conducted. WT n=8 (5 M and 3 F); Tg-SwDI n=9 (6 M and 3 F); Tg+ATZ n=9 (4 M and 5 F); Tg+MTZ n=12 (5 M and 7 F). **D)** Plots for volume for each region. Cortex volume: WT vs. Tg-SwDI, F [1, 15] = 6.629, d = 1.251. **E)** TPM Z-stacks (100 slides per ROI; two ROIs per mouse) of methoxy-positive amyloid deposits (green) were pre-processed with ImageJ (left) and segmented using pixel classification of Ilastik (right). Tg-SwDI, n=28 ROIs in 15 mice (8 F, 6 M); Tg+ATZ, n=21 ROIs in 15 mice (6 F, 8 M); Tg+MTZ, n=24 ROIs in 12 mice (5 F, 7 M). **F)** Plot of Aβ plaque burden (%). **G)** Plot of plaque number per mm^3^. Tg- SwDI vs Tg+MT, LMM, rank-diff = – 15.9 ± 6.31cm, t(35.8) = - 2.513, d = .957). Error bars = SD.

### 2.2 Treatment randomization

For treatment with CAIs, a special diet was designed with either acetazolamide (ATZ; Santa Cruz Biotechnology, Inc. Dallas, Texas, USA) or methazolamide (MTZ; Santa Cruz Biotechnology, Inc. Dallas, Texas, USA). Medication was added at a concentration of 0.01% (100 parts per million), corresponding to 20 mg/kg/day, to a standard diet (Altromin 1320; Brogaarden ApS, Lynge, DK). The three specialized diets were produced with the same pellet form. Researchers involved in the experimental acquisitions were blinded to the treatment. Treated animals were randomly distributed across three treatment groups, each with a specialized diet containing either methazolamide (Tg+MTZ), acetazolamide (Tg+ATZ), or placebo (standard diet – Tg-SwDI), while maintaining balancing between females and males across groups. Since reports have shown that vascular impairments^29^ are already present in the Tg-SwDI mice at the age of 3 months. We anticipated that starting the treatment at 4 months age would slow the vascular impairment in the treated groups. The treatment lasted 6 months, and the scans were performed when the mice were between 10 and 11 months.

### 2.3 Behavioral tests

During the dark phase of the light cycle, mice underwent a behavioral test battery and were acclimated to the testing room for at least 30 minutes prior to each session.

#### 2.3.1 Barnes maze

The Barnes maze assesses visuospatial learning and memory in mice. Our standardized Barnes maze protocol is based numerous past reports using both bright light and a white-noise stimulus to motivate escape from the open, brightly lit maze that is associated with aversive noise^30,31^. The test comprises a raised beige platform (91.5 cm in diameter), with a textured surface, positioned 91.5 cm above the ground, which features 20 evenly spaced holes around the edge, each 5 cm wide, located 2.5 cm from the perimeter. Underneath one of these holes, a gray plastic escape box (10×8.5×4 cm) was placed, and its position was kept consistent throughout trials. The placement of the target hole was varied among mice or groups to ensure balanced spatial conditions. Visual cues were positioned in the testing environment and remained constant across the testing period. Illumination was provided by an overhead lamp (600 lux at the surface of the maze) positioned above the center of the maze. Over 7 consecutive days, mice underwent 12 trials: one habituation trial on day 0, 10 training trials spanning days 1 to 5 (2 trials/day), and the final probe trial on day 6. During the habituation phase, mice were introduced to the maze under an inverted, clear 500 ml beaker for one minute, accompanied by white noise. The white noise was generated by a San Diego Instruments (San Diego, CA, USA) white noise generator located above the maze, with the sound level set at 75dB measured at the maze surface. Subsequently, the beaker was slowly lifted to reveal the target hole, allowing the mouse to explore the escape box for two minutes. Following a 2-hour intertrial interval (ITI) of habituation, standard training trials commenced, with each day featuring two trials; the animal was placed in the center of the maze in a plastic (15×15×20 cm) start box, with a white noise stimulus applied. After 10-15 seconds, the box was lifted, and the animal was given a three-minute window to locate and enter the designated escape hole. Upon successful entry, the white noise ceased, and the mouse was left in the escape box for one minute, and then moved to the home cage. If the mouse failed to find the escape hole within the allotted time, it was gently guided to the target hole beneath the inverted beaker, and left there for one minute, with white noise off. On each training day, trials were separated by 1.5 to 2-hour ITIs, with the maze platform rotated in each trial. On the sixth day, a two-minute probe trial was conducted, like training trials, except that the escape box was removed. Behavior was recorded via an overhead camera for subsequent analysis (Noldus Ethovision XT 11.5 software). Key metrics, including distance traveled and number of non-target holes visited, were measured as indicators of spatial memory and then plotted accordingly.

#### 2.3.2 Open-field test

The open-field test was employed to evaluate animal locomotor activity patterns. The test occurred within a novel open-topped white acrylic enclosure, measuring 40×40×40 cm, with uniform overhead lighting (110-130 lux). Mice were individually introduced into a corner of the arena and given 15 minutes to freely explore. Video of the 30-minute sessions were recorded from an overhead camera, and subsequently analyzed (Noldus Ethovision XT 11.5 software). Locomotor activity parameters, such as total distance traveled, speed, moving time, and time spent in the central zone (20×20 cm) were recorded, and hence plotted.

### 2.4 Surgical preparation and training of animals for optical imaging in the awake state

A chronic cranial window was surgically implanted over the somatosensory area of the barrel cortex (S1bc), as previously described^32^. To minimize stress and enhance recovery, mice were acclimated to handling for 5 days prior to the surgical preparation. On the day of the surgery, mice received a subcutaneous injection of dexamethasone (4.8 mg/kg; Kalcex, Riga, Latvia) to reduce brain post-surgical brain edema. The surgical procedure was performed under anesthesia with 1.75 – 2% Isoflurane (induction with 3%) with 100% O_2_ flow through a face mask. Subsequently, intraperitoneal injections of ampicillin (200 mg/kg) and carprofen (10 mg/Kg) were administered for antibiotic and anti-inflammatory/pain-killer relief purposes, respectively. Lidocaine was applied under the scalp for local anesthesia and eye ointment was applied to both eyes. During the whole surgery, the mouse was securely mounted on a stereotaxic frame (RWD Life Science Co., LTD, Shenzhen, Guangdong, China). A cranial window of ∼3mm in diameter was centered over the region of S1bc representing the whisker C2 (1.5-2 mm anterior-posterior, 3 mm mediolateral). The cranial window was covered with an in-house glued glass plug consisting of one 5-mm and three 3-mm cover glasses. The glass plug was secured to the skull applying cyanoacrylate glue around the edges (Loctite Super Glue gel, Loctite® Brand, Westlake, Ohio, USA). A custom-made head-bar was affixed and glued onto the frontal area of the skull and the whole exposed area of the skull was covered with a layer of dental cement (Meliodent, Kulzer. Hanau, Germany). After surgery, mice were recovered in a heated recovery chamber and were closely monitored until voluntary movement was observed. Subsequently, each mouse was moved back to its own housing box. We continued to monitor their recovery for a period of 5 days post-surgery, providing antibiotic and anti-inflammatory medication as needed. Acclimatization with handling continued through the post-surgical period to minimize stress and improve training sessions.

After the post-surgical recovery period, mice underwent daily training sessions to acclimatize them to awake imaging conditions, with training durations progressively increasing from 15 minutes to the target session length of ∼2.5 hours over a minimum of 10 days. This protocol was designed to minimize stress levels, as demonstrated by prior research from our group showing significant reductions in corticosterone levels and heart rate in mice after 10–11 days of such training, indicating stress habituation for both male and female mice^33^. For training sessions, we used a replica of the custom-built imaging frame to ensure familiarization with the restraint apparatus. After each session, mice were rewarded with condensed milk and returned to their cage. During the scanning session, mice were monitored through a night vision camera installed within the imaging systems. A rest day was given between imaging sessions. The gradual habituation process implemented in our study was specifically designed to reduce stress-induced vascular perturbations based on our previous experience^33,34^, thereby minimizing confounding effects on hemodynamic measurements. However, if hyperreactivity to handling or restrain was excessive, the scanning session was delayed after two additional days of training.

We placed a tail catheter for infusion of fluorescent dyes to label the plasma for TPM imaging. For this procedure mice were lightly anesthetized with Isoflurane 1.2% delivered through a face mask (100% O_2_ flow). A catheter was assembled with a PE10 tube attached to a 50-unit (500 µl) insulin syringe on one side and a 30G Monoject needle (Covident) on the opposite side. The catheter was filled with a 5% heparin (LEOPharma, Ballerup, Denmark) solution. The catheter was placed on one of the lateral tail veins, after warming up the tail and applying ethanol to induce vasodilation. The catheter was fixed with dental cement (GC-Fuji-Plus, GC Corporation, Japan) to avoid movement or detachment during the scanning period. After the scanning session, the catheter was gently removed, and the tail was gently pressed until bleeding stopped.

### 2.5 Optical imaging acquisition

We performed imaging in awake head-restrained mice using a frame equipped with a head-holder and a fabric cradle where the mouse could rest in a relaxed position. optical coherence tomography (OCT). For OCT imaging, we applied a spectral-domain OCT system (1310 nm center wavelength, bandwidth 1700 nm, Telesto II, Thorlabs, Inc. Newton, NJ, USA). This system provides an axial resolution of ∼3.5 um and an imaging speed of ∼47,000 A-scan/s. To achieve a transverse resolution of ∼1.5 µm, we used a 10X near-infrared (NIR) objective (Mitutoyo Corporation, Kawasaki, Kanawaga, Japan). We acquired time-series OCT-angiograms, each consisting of 400 x 400 pixels (600 µm^2^), using the decorrelation method^35^. In our setup, each OCT-angiogram acquisition took ∼8.5 s with a buffering time of 5 s between OCT volumes. Sixty volumes of the cortical microvasculature were acquired in each region-of-interest (ROI).

#### 2.5.1 Detection of capillary stalls from post-processed data

After postprocessing the OCT-angiograms, a volume of 30 pixels was selected 135 - 225 µm beneath the brain surface and was projected as maximum intensity projection (MIP) for analysis. In OCT imaging, stalling events are considered capillaries without perfusion during an OCTA volume. OCT signal relies on the movement of particles and light path interference. Each volume represents ∼8.5 seconds of capillary dynamics. Stalled capillaries typically re-perfuse within one or two volumes, although some stalls may last longer. We identified capillary stalling events using the graphical user interface (GUI) ‘CapStall’ designed in MATLAB (Mathworks) that allowed a semiautomatic identification of the stall events^36^. For postprocessing, two research assistants, blinded to the treatment randomization, inspected all images in a time series (volume scanned as a function of time) to identify all image frames in which the capillary segments stalled at least one time. After having done the identification of stalling capillaries, a stall-o-gram was plotted. Stall-o-grams depicted the capillary stalls and the stalling frequency during the 60 (1 volume = 8.5 seconds) volumes scanned. For statistical analysis, the assistants manually quantified the capillaries examined in each MIP using Image J and a MATLAB script to calculate the following parameters:

a. Total of identified capillary stalls.
b. Incidence rate, defined as the fraction of capillaries showing a stall episode for 8.5 minutes (sixty 8.5-second volumes).
c. Point of prevalence, which is the average number of capillary stalling episodes per volume (8.5 seconds).
d. Cumulative stall time, representing the cumulative fraction of stalling time relative to the total scanning time (∼8.5 minutes).

#### 2.5.2 Two-photon imaging

All two-photon microscopy (TPM) imaging experiments were performed using a Bruker Ultima-IV two-photon laser scanning microscope (Brucker Corporation, Billerica, MA, USA) and Praire View Software version 5.1 (Brucker Corporation, Billerica, MA, USA). Excitation of fluorescent dyes was performed with a tunable Ti:Saphire laser Chameleon (Coherent, Santa Clara, California, USA). Fluorescence emission was detected by two GaAsP photomultiplier modules (Hamamatsu, H10770. Hamamatsu City, Japan). We used an Olympus 10x water immersion objective (0.3 numerical aperture – NA), 3.3 working distance – WD) to image a 1.18 mm^2^ FOV for bolus tracking, intravascular pO2 estimations, and reference angiograms for bolus tracking. We used an Olympus 20x water immersion objective (1.0 NA, 3.00 mm WD) for capillary line-scans, high resolution z- stacks for angiograms, and Aβ plaques. To avoid any effect of temperature on brain hemodynamics, we heated the objective lens with an electric heater (TC-HLS-05, Bioscience Tools) to maintain the temperature of the water between the cranial window and the objective lens at 36–37°C.

Two imaging sessions were performed, interleaving one day between scans to reduce stress induced by repetitive sessions. All scans were performed during steady-state conditions. On the first day of TPM imaging, we acquired measurements of brain hemodynamics in awake-restrained mice. The examinations on day two did not include data that are sensitive to movement and physiological variations (e.g., capillary diameter), and they were therefore performed under slight anesthesia (0.9 – 1% Isoflurane) delivered through a face mask as we acquired morphological data to avoid excessive movement during the ∼10 min acquisition of each z-stack (i.e., angiograms, Aβ plaques).

TPM acquisition sequences:

1. Indicator-dilution technique (Bolus tracking): We estimated MTT and CTH applying an indicator-dilution methodology as previously described^37^. This methodology relies on the detection of concentration-time-curves (CTC) during the passage of a bolus of fluorescent dye from the arterial to the venous network. Each bolus was injected through the tail catheter using an infusion pump (GenieTouch; KenScientific, Torrington, USA) set at an infusion rate of 20 µL/s. To avoid pain due to the I.V. infusion of cold solutions, we kept the catheter warm with a rubber tube attached to a standard heating circulation bath (ThermoScientific, Walthman, USA). To locate the ROI, we identified the shadows of the pial vessels detecting NADH- autofluorescence (820 nm excitation). The autofluorescence signal was filtered with a 460 ± 25 nm bandpass filter (Chroma Technology, USA). The volume of the catheter was ∼60 µl, and was filled with heparin solution before the scan; therefore, we habituated the mice to the bolus injection ∼6 times (10 µl boluses) before performing the recording paradigm to fill up the catheter with the fluorescent dye. We identified arterial and venous networks during the habituation paradigm. Each bolus consisted of 10 µl of the fluorescent dye Texas-Red dextran (70,000 M.W.; Thermo Fischer Scientific, USA). The signal was filtered with a 605 ± 35 nm bandpass filter (Chroma Technology, USA). We identified an ROI as an area where both pial arteries and veins converged, and a large region with capillaries was observed. Each bolus was infused after 7 seconds from the initiation of a time-series of 2D spiral scans with a spatial resolution of 256 x 256 (pixel size = 2.30 µm) which yielded a frame rate of ∼6.98 frames-per-second (fps). A total of 3 boluses of each mouse were included for analysis.
2. Intravascular pO_2_ measurements: After the placement of the tail catheter, a bolus of ∼40–50 µl (10–15 µM final concentration in the blood plasma) of the phosphorescent probe PtP-C343 (MW ∼32,000)^38^. Using the residual green fluorescence of the C343-antenna fragments in PtP-C343, we performed a raster scan to map the pial vessels and quantify their diameter. The fluorescent signal was detected with a PTM and passed through a 525 ± 25 nm bandpass filter. We used the PointScan mode of the TPM software to obtain single-point measurements within the luminal side of the arterial and venous networks of the most superficial layers of the brain cortex. We aimed to include two points per vessel in the same vessel detected during the indicator-dilution scans. Upon excitation at a single point during 10 µs (820 nm, ∼10 mW average power with beam modulation off), the phosphorescence output was passed through a 680 ± 30 nm bandpass filter and collected during 290 µs by a PMT module (H10770PA-50; Hamamatsu), as previously described^39^. For O_2_ estimates, we averaged ∼15,000 phosphorescence decays per point. Quenching of the probe’s phosphorescence is dependent on the O_2_ concentration. Therefore, the phosphorescence lifetimes in arteries are expected to be shorter than in veins. Two trials per mouse were performed.
3. Line-scans for capillary scans: After the indicator-dilution scans, the remaining volume required to reach 200 µl was injected through the tail catheter to enhance contrast for capillary scans. Capillary scans were conducted using Linescan mode of the TPM software with a 20x featuring a 1.0 N.A. and W.D. of 2.0 mm (UMPLFLN 20x, Olympus). Digital zoom was set to 5x, resulting in an *XY* pixel size of 0.23 µm. Capillaries were randomly selected, with efforts made to remain within the same FOV as used for indicator-dilution scans. Fifty to twenty capillaries were scanned per mouse through depths between 50 – 200 µm from the surface (Z = 0). The number of capillaries was chosen to balance the time constraints of the scanning session, which also included additional methodologies. Depth was selected based on previous observation of low variability in cell flux and velocity within this range^37,40^. A scan path was assigned in 1 – 5 capillaries, once in the axial direction of the capillary lumen for RBCv estimation, and twice transversally to the lumen for an averaged cell flux and capillary diameter. Line scan period varied between 0.5 to 2 ms per line. Scans were performed under steady-state conditions for 5 seconds.
4. Z-stacks for Aβ detection and angiograms: For detection of Aβ plaques, the day before to the scanning session, the mice were injected via I.P. with a dosage of 2 mg/kg of Methoxy-X04 dye (Tocris Bioscience, Bristol, UK). In our setup, the excitation peak was at 790 nm, which was selected for the scanning sessions. The signal was filtered with a 460 ± 25 nm bandpass filter. For the detection of vessels, we labeled plasma with 200 µl of Texas-Red Dextran 70,000 MW (5 mg/ml, Thermo Fischer Scientific, Waltham, MA, USA). Excitation of Texas-Red Dextran was performed using a wavelength of 900 nm and emission was filtered with a 605 ± 35 bandpass filter.

#### 2.5.3 Post-processing of indicator dilution techniques

MTT and CTH were estimated by modeling the tissue transport function (TF) through deconvolution following the injection of a dye bolus, as previously described^37^. For the analysis of the 2D time-series, we developed a dedicated MATLAB GUI that allowed the selection of vessels within the FOV. For modeling TF, it was crucial to select an arterial input function (AIF) and a venous output function (VOF). Initially, we identified pial arteries and veins as vessels positioned parallel to the cortex and without diving into the brain tissue. The main feeding arterial AIF was selected as the artery with the largest diameter and the shortest time-to-peak (TTP) of fluorescence intensity. Venous VOF was defined as the vein with the largest diameter and with the longest TTP. We identified the main arteriolar AIF as a vessel branching from an artery (10 < *d* < 30 µm), with the shortest TTP. The main venular VOF was defined as an ascending vessel (10 < *d* < 60 µm) with an anastomosis with a pial vein and with the shortest TTP. The vascular TF described the relative amount of dye emerging at the VOF as a function of time after an idealized, instantaneous bolus injection on the AIF. The TF was modeled by a gamma probability density function with parameters α and β, and a scaling factor, f. With this TF, we estimated MTT as the mean (α·β) and CTH as the standard deviation (√α·β). To visualize curve fitting and verify the MTT estimation, the AIF and VOF were compared after time-shifting the VOF curve by the estimated MTT (Supplementary Fig. 1). Bad curve fitting or curve-intensity saturation were used as criteria for rejection of analyzed pairs (Supplementary Fig. 2).

#### 2.5.4 Post-processing of lifetime imaging for pO_2_ measurements

To estimate pO_2_, we determined the phosphorescence lifetime by fitting the phosphorescence intensity decay with a single-exponential function using the nonlinear least-squares method. The lifetime was converted to pO_2_ using the calibration plot obtained in independent oxygen titration experiments^38^. For statistical analysis, we averaged the two measurements per vessel. To correlate oxygen tension to the diameter of the pial vasculature, segmentation, and diameter estimation were performed by two research fellows, blinded to the treatment, averaging the distance of 5 transversal lines placed transversal to the vessel axis using the ROI manager plug-in of FIJI.

#### 2.5.5 Post-processing of single capillary scans

Before processing, all images were filtered using a top-hat filter to improve SNR. RBCv, capillary cell flux, and diameter were calculated using a MATLAB GUI, as previously performed^37^. Briefly, RBCv was calculated from the axial capillary line scan using the Radon transform algorithm for for accurately and robustly analyzing streak angles^41^. Calculations were performed in 150 ms windows, displaced by 50 ms for each iteration. Cell flux was calculated by analyzing the intensity variations from the cross-section scan of each capillary. Average intensity profiles were derived from 150-ms time interval windows within transversal line scans. A cluster analysis was applied to determine the presence of cells as they passed through the capillary. The capillary diameter was estimated from cross-sectional scans as the full width at half-maximum (FWHM) of intensity profiles within 150 ms windows^42^, disregarding time intervals with the presence of RBCs.

#### 2.5.6 Post-processing of angiograms for capillary morphometrics

The identification and segmentation of the capillary network involved the following steps:

1. Raw image stacks of the vasculature were first pre-processed using a Macro written in ImageJ (National Institutes of Health, Bethesda, Maryland, USA) for contrast enhancement and image denoising. The macro involved the following steps:

a. CLIJ2_equalizeMeanIntensitiesOfSlices for bleaching correction.
b. Contrast enhancement with normalization (saturated pixels = 0.4%).
c. Background subtraction (rolling ball = 50).
d. 3D Filtering (Median 3D; x = 2, y =2, z=2).
2. Vessel segmentation was performed using DeepVess, a deep convolutional neural network that enables the identification of individual capillary segments by extracting the vessel centerline^43^.
3. Identification of capillary segments after segmentation was performed using a dedicated MATLAB script previously described^43^. Postprocessing included filtering according to diameter to remove the large vessels, with a set diameter limit of 8 pixels, equivalent to 9.2 µm (*XY* resolution = 1.15 µm per pixel). The vessel diameter and its standard deviation (SD) were calculated for each segment by measuring the distances of all non-zero voxels within the segment’s binary skeleton to the nearest boundary of the vessel segmentation. The diameter was defined as twice the mean of these distances, while the SD was computed from the same distance measurements to capture the variability in vessel diameter along each segment.
4. Finally, another dedicated MATLAB script was employed to estimate capillary tortuosity as the capillary segment length divided by the Euclidean distance between the segment endpoints.

Additional parameters estimated: 1) Capillary diameter coefficient of variance as diameter standard deviation along the segment divided by the mean diameter along the segment; 2) Capillary length density as the total capillary length (µm) in the total z-stack volume (µl); and 2) Capillary blood volume as the fraction of the sum of capillary volumes (µl) and the total z-stack volume (µl).

#### 2.5.7 Post-processing for Aβ deposits quantification

Aβ quantification was performed using a combination of Macros designed using ImageJ and Ilastik, an interactive machine learning for bioimage analysis^44^. A sub-stack of 100 slices (100 µm) was selected from a z-stack scanned with TPM to keep consistency between all groups. Contrast enhancement and denoising were performed using the same Macro as described for the angiograms. Segmentation of plaques was conducted using the pixel classification workflow of Ilastik, which assigns labels based on pixel features and user annotations using a Random Forest classifier. The software was trained using two sub-stacks and the rest of the stacks were classified using the batch mode. The Ilastik model’s performance was validated using the raw binary image as the ground truth, as it represents the amyloid signal in its entirety, free from noise and additional artifacts. The Ilastik model’s performance was validated using the raw binary image as the ground truth, as it represents the amyloid signal in its entirety, free from noise and additional artifacts. The model’s segmentation accuracy was assessed on 25- slice and 100-slice test sets to evaluate its effectiveness on three-dimensional image stacks. For the 25-slice test set, the model achieved a Dice Similarity Coefficient (DSC) of 0.832, Precision of 0.801, and Recall of 0.866. On the 100-slice test set, the model demonstrated improved performance, achieving a DSC of 0.843, Precision of 0.766, and Recall of 0.938. These results indicate robust segmentation performance, particularly in capturing true positives (high Recall), and a consistent overlap between the predicted and ground truth segmentations (DSC). Finally, for quantification of Aβ deposits, we use the ‘3D Objects Counter’ from ImageJ, which gave the volume of each plaque across the sub-stack.

### 2.6 Extraction of tissue samples for molecular analyses

Following the last imaging session, mice underwent euthanasia by induction of deep anesthesia with isoflurane (4% in FiO_2_ 25%) and subsequent administration of an overdose of pentobarbital (100 mg/kg) intraperitoneally. Upon confirmation of respiratory arrest, a transcardial extraction of blood (300 – 500 µl) was performed. Plasma was recovered after centrifugation to remove the blood cells. Subsequently, the brain was carefully extracted from the skull of each mouse. The hippocampus and the region of the cortex of the scanned area were extracted (left hemisphere). All tissue samples were promptly snap-frozen and stored at –80°C until further preparation for molecular analyses.

### 2.7 Enzyme-linked immunosorbent assay (ELISA) to determine specific protein levels

The frozen tissue samples were finely minced on an aluminium plate cooled below 0 °C, and further homogenized by grinding in a 1.5 mL LoBind (Eppendorf) tube with a plastic pestle (GE Healthcare, Sample Grinding Kit), in 20 μL pr. mg tissue of an extraction buffer with 2% SDS, 100 mM sodium HEPES, and 1 cOmplete™ Protease Inhibitor Tablet (Roche) pr. 10 mL (1x PI). Finally, the sample was ultrasonicated for three cycles of five ultrasonication pulses on a Branson Sonifier 250 (Branson Ultrasonics Corporation) at output control 3 and 30% duty cycle. The protein content of each sample was measured in triplicates with the Pierce™ BCA Protein Assay Kit (Thermo Scientific) on a BSA standard curve according to the manufacturer’s protocol.

Samples were diluted in extraction buffer to 2 μg protein/μL. To replace SDS with the ELISA compatible buffer Tween-20, SDS was precipitated was by addition of an equal volume of a Tween/K buffer with 0.1% Tween-20, 100 mM K2HPO_4_, 1×PI buffer followed by centrifugation for 15 minutes at 15,000×g at 2 °C, to pellet the potassium-SDS precipitate. The supernatant was aliquoted for each of the ELISAs, and total protein content was determined, in triplicate, with the BCA assay.

For use in the ELISA standard curves, a sample buffer was prepared by precipitating SDS from the extraction buffer with the Tween/K buffer as described above. All samples and standards were measured in duplicate. Absorbance was measured at 450 nm. The observed absorbance in the 0 standard, was subtracted from all measurements. A 4-parametric standard curve was applied to calculate specific protein concentrations. The average of the duplicates was multiplied by the dilution factor and normalized to the total protein concentration of the sample after SDS-removal. The results are reported as pg-specific protein/mg total protein.

Aβ-40 was detected by an ELISA kit (Invitrogen, Cat. #: KHB3481) targeting human Aβ-40. Transgenic mice samples were diluted by a factor of 200 in the kit ‘Standard Diluent Buffer’. The Aβ-40 standard was reconstituted in 55mM sodium bicarbonate, pH 9.0, according to protocol, and diluted from 100 ng/mL to the range of the standard curve (7.81-500 pg/mL) in a 1:199 mix of sample buffer and kit ‘Standard Diluent Buffer’. The assay was otherwise run according to the manufacturer’s protocol. The manufacturer reports a 0.5% cross-reactivity with rodent Aβ-40 which may explain the trace amount signal observed in some samples from WT mice, which were furthermore diluted only by a factor of 5.

For the detection of Aβ-42, we used an ELISA kit (Invitrogen, Cat. #: KHB3441) targeting human Aβ-42was employed. Samples were diluted by a factor of 5 in the kit ‘Standard Diluent Buffer’. The Aβ-42standard was reconstituted and diluted in a 1:4 mix of sample buffer and kit ‘Standard Diluent Buffer’. The assay was run otherwise according to the manufacturer’s protocol.

For detection of CypA we used an ELISA kit (Abbexa, Cat. #: abx585050) targeting murine CypA. Samples were diluted by a factor ca 1.3 in the kit ‘Standard Diluent Buffer’ (80 μL sample + 25 μL kit buffer). The CypA standard was reconstituted and diluted in a 3.8:1.2 mix of sample buffer and kit ‘Standard Diluent Buffer’. Color development was allowed to proceed for 60 min. The assay was run otherwise according to the manufacturer’s protocol.

ICAM-1 was detected using an ELISA kit (Abcam, Cat. #: ab100688) targeting murine ICAM1. Samples were diluted by a factor of 3.6 in the kit ‘1x Assay Diluent B’ (28.5 μL sample + 103.5 μL kit buffer). The ICAM1 standard was reconstituted and diluted in a 30:11.4 mix of sample buffer and kit ‘Standard Diluent Buffer’. The assay was run otherwise according to the manufacturer’s protocol.

### 2.8 Mesoscale Discovery

Quantification of VEGF-A in plasma samples (1:1) obtained from scanned animals was conducted using the U-Plex Mouse VEGF-A Mesoscale Assay (Mesoscale Discovery, Rockville, USA) based on electrochemiluminescence. The procedure adhered to the standard protocol recommended by the manufacturer for preparation and measurements.

### 2.9 Ex-vivo MRI

An independent sample of mice was used for ex-vivo MRI scans. Four experimental groups were studied: WT mice (C57BL/6J, N=8 (F = 3, M = 5)], Tg-SwDI mice treated with MTZ (Tg+MTZ, N=12 (F = 7, M = 5)], Tg-SwDI mice treated with ATZ (Tg+ATZ, N=9 (F = 5, M = 4)] and Tg-SwDI mice treated with placebo (Tg-SwDI, N=9 (F = 3, M = 5). Ex-vivo MRI was chosen over imaging in anesthetized mice due to logistical constraints in monitoring the animals during extended scanning sessions.

Upon euthanasia, mice underwent perfusion-fixation with heparin (10 U·mL−1)-treated 0.9% saline (pH = 7.3) for 2 minutes, followed by ice-cold 4% buffered paraformaldehyde (PFA, pH = 7.2–7.4) for 2 min. Subsequently, the brain was removed from the skull and placed in PFA for extended fixation. In preparation for imaging, samples were washed in phosphate-buffered saline (PBS, Sigma USA, P4417- 50TAB) for a minimum of 24 hours to improve the signal by removal of excess fixative. The sample was then securely placed in an MRI-suitable tube filled with fluorinert (FC-770, 3M inc.) as is standard^45^.

MRI was conducted using a Bruker Biospec 9.4T preclinical MRI system (Bruker Biospin, Ettlingen, Germany) equipped with a bore-mounted 15 mm volume coil. High-resolution structural data for volumetric estimation was acquired at 150 μm in-plane resolution in 200μm thick slices using a segmented (8 segments) T2-weighted echo planar imaging (EPI) sequence. Scan parameters were 16 avs, TE=31.7 ms, TR = 4500 ms, BW= 277 kHz, 70 axial slices. ROIs (i.e., cortex, somatosensory cortex (SMS), hippocampus (HC), and white matter WM) were manually segmented using ITK-SNAP, as shown previously^46^. The volume of each of these regions was then calculated as the number of voxels in each ROI multiplied by the nominal voxel volume (150µm×150 µm×200µm, i.e. 0.0045 mm^3^ pr voxel).

### 2.10 Statistical analysis

We used R version 4.0.3 to perform statistical to assess our two hypotheses: (1) baseline model differences (WT vs. Tg-SwDI), and (2) treatment effectiveness, comparing Tg+ATZ, Tg+MTZ groups relative to Tg-SwDI. For behavioral analysis and MRI analyses, a two-way an analysis of variance (ANOVA) was performed to determine overall differences among the groups (WT, Tg-SwDI, Tg+ATZ, Tg+MTZ) with sex (female, male) as a between-mouse factor.

Parametric analysis was performed for normally distributed data (evaluated using Q-Q plots, Shapiro-Wilk tests, or the Kolmogorov-Smirnov test for larger datasets). For variables that did not meet normality assumptions, we applied log-transformation and reassessed normality. If normality was achieved following log-transformation, parametric analyses were performed as described below. For variables that remained non-normal after log-transformation and without repeated measurements per mouse, we conducted non-parametric analyses using either the Kruskal-Walli’s test and pairwise Wilcoxon tests with Benjamini-Hochberg correction.

In all analyses of optical imaging data, outliers were identified by calculating the z-score of each variable and removing data with a z-score below –3.29 or above 3.29, as previously described^47^. We used the lme4 package for linear mixed-effect model (LMM)^48^ estimates and constructed models with the hemodynamic parameter (e.g., cumulative stalling time, MTT, CTH) as dependent parameter. As independent parameters, we included group category as a fixed effect and subjects as a random effect (e.g., *MTT ∼ group + (1 | mouse ID)*). For variables that did not meet normality assumptions, log-transformed values were used after testing normality again. For variables that remained non-normal after log-transformation, we conducted LMM with ranked values for semi-parametric analyses to account for the repeated measurements.

To examine the effect of gender, we further expanded the model to include gender as an additional fixed effect. These two models were compared using a likelihood ratio test using the function anova in R (e.g*., anova(model with sex effect, model without sex effect)*) to assess whether including sex as a fixed effect significantly improved the model fit. If a significant effect of sex was observed, the parameter estimates were adjusted for sex in subsequent analyses.

If the LMM encountered a singular fit, indicating that the random intercept for mouse did not add value, we simplified the model to a linear model (LM) without the random effects. Multiple comparison was performed using estimated marginal means (EMMs) with the ‘emmeans’ package based on either the LMM or the simplified LM. EMMs were calculated for each group, and specific contrast were defined to evaluate the baseline model differences and treatment effectiveness. Custom contrasts were applied with BH adjustment to control for multiple comparisons.

We estimated effect sizes by calculating Cohen’s d^49^ for parametric analysis reaching significance or trends (p ≥ .05 and < 1). Effect sizes were categorized as small (d=.2), medium (d=.5), and large (d≥.8) based on standard conventions. For non-significant trends, power analysis to determine the sample size required to achieve 80% power (1-β = .8) at a significance level of α=.05. This approach rigorously evaluates trends, providing insights into both their statistical significance and biological relevance.

To assess the relationship between MTT and CTH, we performed a correlation analysis across two parts of the vascular networks: artery-to-vein and arteriole-to-venule pairs: artery-to-vein and arteriole-to-venule pairs. This analysis utilized a linear model (LM) to determine the strength and direction of correlation within each group. The models were adjusted by the bolus trial to control for variability introduced by bolus injection characteristics.

All normality tests and the statistical test or models selected for each variable are shown in Supplementary Materials 1.

All plots were constructed using the ‘ggplot2’ package, displaying individual data points along with error bars representing the standard error of the mean (SEM). The results in the tables are presented as mean ± S.D.

## 3. Results

### 3.1 Middle-aged Tg-SwDI mice are characterized by normal memory function, despite amyloidosis and motor function abnormalities

To confirm that 9–10-month-old Tg-SwDI mice are a suitable model of presymptomatic AD, we compared spatial memory and locomotion to those of wild-type (WT) mice, using Barnes maze. Notably, no significant changes in the number of errors or latency were observed between WT and Tg-SwDI mice in the Barnes maze (Fig. 1A), demonstrating preserved spatial memory function in the current settings. Our assessment was coupled to open field test to assess locomotor activity, given that escape latency in the Barnes maze may be influenced by non-cognitive factors such as motor function, motivation, anxiety or general search strategy^50,51^. The test revealed motor changers in the Tg-SwDI mice, characterized by a reduction in the distance travelled compared to WT, accompanied by a trend toward reduced moving time (ANOVA, p = .0654), with a moderate effect (Cohen’s d = .746, Fig. 1B). An estimated sample size of 30 mice per group required to achieve statistical significance. These results demonstrate that in our settings, the changes in motor function, did not influence the overall latency in Barnes maze. This is consistent with recent studies that found no spatial learning and memory deficits in 9-month-old Tg-SwDI mice^27,28,52^.

To detect structural, cortical, or hippocampal changes, we estimated regional tissue volumes by ex-vivo MRI in a group of 9-11 months old Tg-SwDI and WT mice. Cortex, hippocampus (HC), somatosensory cortex (SMS), and white matter (WM) were segmented, providing volumes for each region (Fig. 1C). Notably, Tg-SwDI mice did not show signs of atrophy but rather a trend towards increased cortical volume compared to the WT (Kruskal-Wallis, p = .0833, Fig. 1D). The observed trend corresponds to a large effect size (Cohen’s d =1.121). These results are consistent with previous observations of whole-brain hypertrophy in another model of cerebral amyloidosis^53^ and observation in AD patients^54^.

To detect Aβ burden, we used volumetric two-photon microscopy (TPM) scans and the Aβ fluorescent tracer, methoxy-X04 (Fig. 1E). We confirmed that 9-10 months old Tg-SwDI mice show extensive Aβ burden at 9 to 10 months old (Fig. 1F – 1G).

Overall, our analysis of the Tg-SwDI strain in the 9–11-month age range revealed no brain atrophy or no deficit in spatial learning and memory despite the presence of Aβ in the brain tissue, underscoring its suitability as a model for the presymptomatic period of the AD continuum.

### 3.2 Capillary flow disturbances are present in presymptomatic Tg-SwDI mice

We examined presymptomatic Tg-SwDI mice for capillary stalling, a phenomenon first described in AD models and linked to reduced CBF^15^. Using optical coherence tomography angiography (OCT-A; Fig. 2A – 2D)^36^, we observed an increased number of capillary stalls in Tg-SwDI mice compared to WT mice (Fig. 2E and Supplementary Table 1), including higher incidence (Fig. 2F) and prevalence (Fig. 2G). However, no significant difference in the cumulative stall duration was found between WT and Tg-SwDI mice (Fig. 2H). We did not observed correlation between the incidence of stalling events and the cumulative stall duration (Supplementary Fig. 3B), indicating that the frequency of stalling events may not predict the total time capillaries remain stalled.

**Figure 2.**
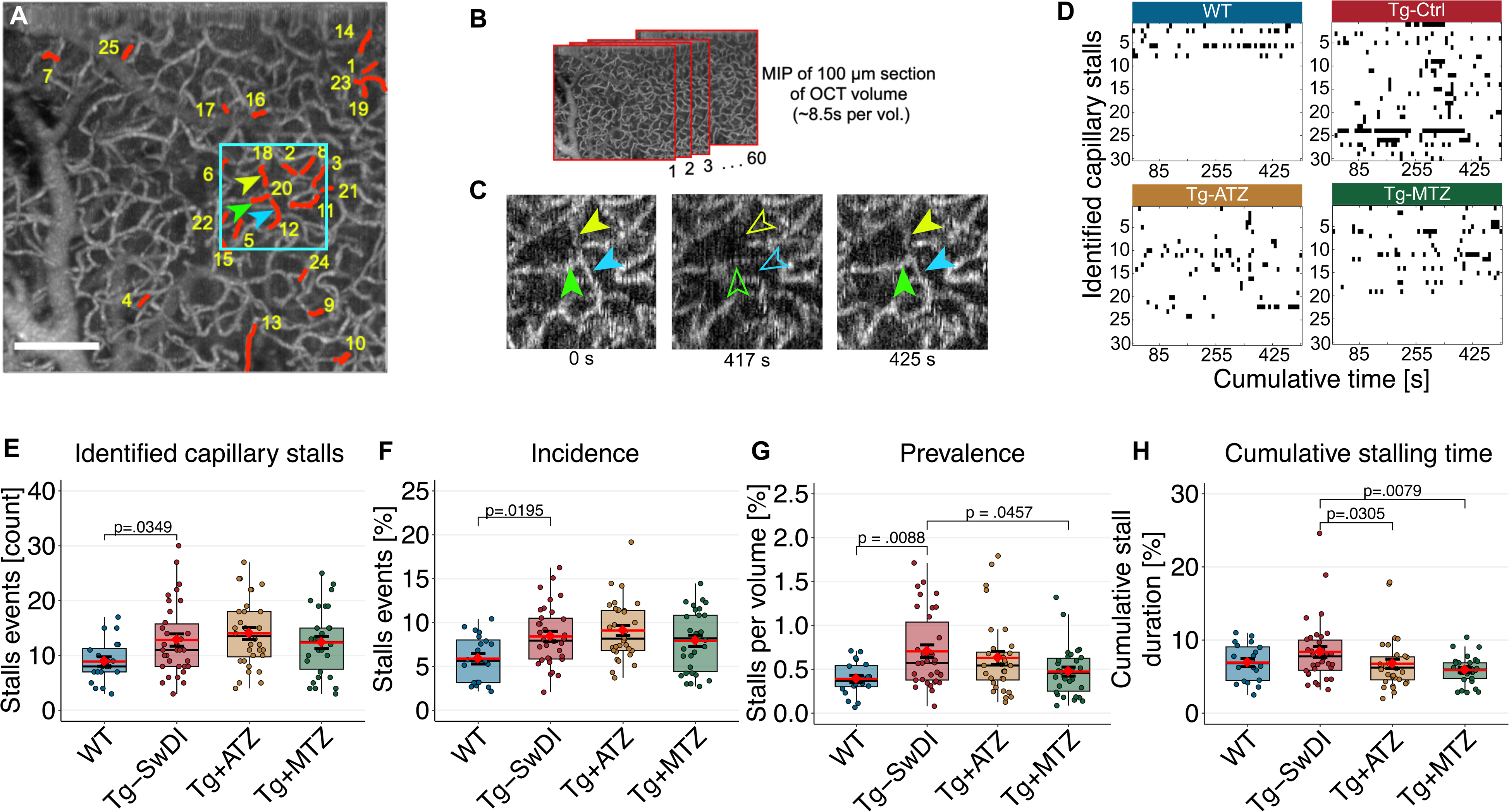
Long-lasting capillary stalling in the Tg-SwDI is prevented by CAIs. **A)** OCT-A was used to detect stalling events (ROIs per mouse). WT n=20 ROIs in 11 mice (6 F, 5 M); Tg-SwDI n=34 ROIs in 18 mice (11 F, 7 M); Tg+ATZ n=32 ROIs in 17 mice (7 F, 10 M); Tg+MTZ n=30 ROIs in 15 mice (7 F, 8 M). MIP of 100 µm (axial) of an ROI (600 µm^2^) with stalling segments indicated (red markings). **B)** Sixty OCT-A-volumes in 2 ROIs per mouse were performed (∼8.5 seconds/volume). **C)** Magnification of selected region (cyan in a) with capillary segments (arrowheads) with temporary perfusion interruptions. Hollow arrowheads indicate a stalling event. **D)** Representative stall-o-grams from a ROI per group of stalling segments through 60 OCT-A-volumes. Black points represent stalling events per each image frame. **E)** Identified capillary stall events in each examined ROI. WT vs. Tg-SwDI, LMM, log-scale difference (log-diff.) = 4 ± 1 stalls, t(63.1) = 2.398, d = .721. **F)** Incidence of capillary stalls. WT vs. Tg-SwDI, LMM, diff. = 2.52 ± 1.05 %, t(63.1) = 2.398, d = 1.002. **G)** Prevalence as a percentage of stalling events per volume. For comparisons between Tg-SwDI and WT group, a ranked LMM was used to account for non-normality. WT vs. Tg-SwDI, LMM, rank-diff. = 26.1 ± 9.63, t(60.4) = 2.707, d = .882; Tg-SwDI vs. Tg+MTZ, LMM, log-diff. = –.389 ± .167, t(58.1) = −2.338, d = .657. **H).** Cumulative stall duration as percentage of time of each ROI (∼8.5 minutes). Tg-SwDI vs. Tg+ATZ, LMM, log-diff. = –.224 ± .102, t(116) = −2.190, d = .539; Tg-SwDI vs. Tg+MTZ, log-diff. = −.306 ± .104, t(116) = –2.941, d = .737. Scale bar = 100 µm. Error bars = SD

We also examined single capillary hemodynamics using TPM (Fig. 3A). We found lower red blood cell velocities (RBCv; Fig. 3B) and cell flux (Fig. 3B and Supplementary Table 2) in Tg-SwDI mice compared to WT mice. Capillary linear density (flux/RBCv), and diameter were similar across the two groups (Fig. 3D – 3E), suggesting capillary perfusion is maintained, consistent with previous observations in 12-month-old Tg-SwDI mice^55^. However, the reduced RBCv may indicate subtle impairments in flow regulation, potentially reflecting early capillary dysfunction or altered vascular tone in presymptomatic Tg-SwDI mice.

**Figure 3.**
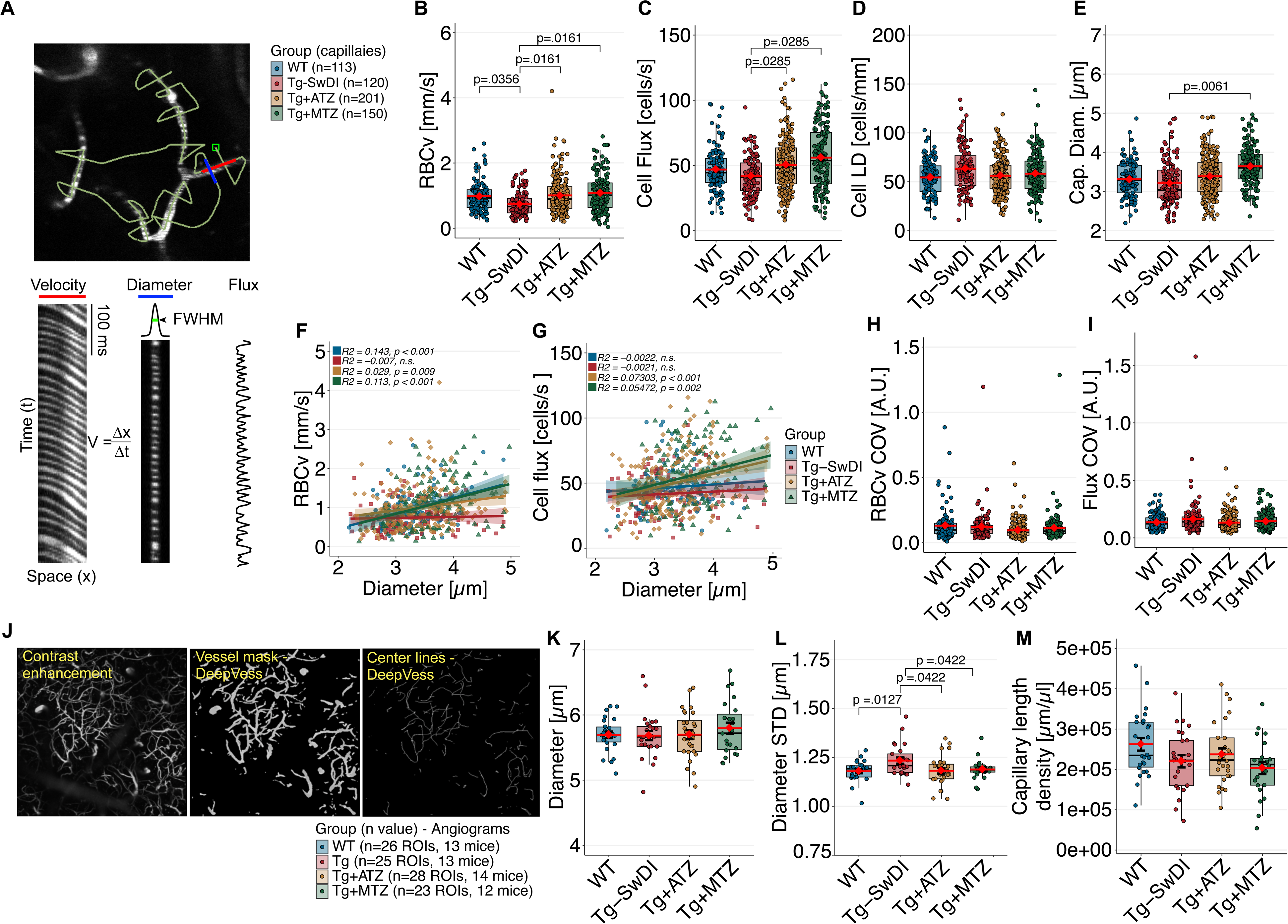
Disturbances in capillary flows and morphology are prevented in CAI- treated Tg-SwDI mice. **A)** Simultaneous estimation of RBCv, cell flux, capillary LD and capillary diameter (Cap. Diam.) from single-capillary scans. A path traversed the capillary longitudinally (red) and transversely (blue) were TPM linescans were performed (upper left). RBCv (*bottom left*) was calculated using the Radon transformation based on streak angles^41^. Diameter (*bottom middle*) and cell flux (bottom *left*) were estimated from the transversal. A total of 583 capillaries were scanned across groups (WT, 6 mice (3f, 3m); Tg-SwDI, 7 mice (5f, 2m); Tg-ATZ, n = 10 mice (5 F, 5 M); Tg-MTZ, n = 9 mice (6f, 3m). Scale bar: 20 µm. **B)** Plot of the RBCv. WT vs. Tg-SwDI, LMM, log-diff = –.282 ± .128, t(31) = –2.197, d = .548. Tg-SwDI vs. treated groups, ranked LMM to account for non-normality. Tg-SwDI vs. Tg+ATZ, rank-diff = 92.2 ± 35.9, t(31.7) = 2.568, d = .608. Tg-SwDI vs. Tg+MTZ, rank-diff = 94.5 ± 37.3, t(33.0) = 2.537, d = .624. **C)** Plot of the capillary cell flux. For comparisons a ranked LMM was used to account for non-normality. Tg-SwDI vs. Tg+ATZ, rank-diff = 72.9 ± 34.5, t(31.5) = 2.296, d = .515. Tg-SwDI vs. Tg+MTZ, rank-diff = 91.4 ± 35.8, t(32.9) = 2.549, d = .594. **D)** Plot for capillary linear density (Cell flux/RBCv). **E)** Plot of capillary diameter. Tg-SwDI vs. Tg-MTZ, LMM, log-diff = .1087 ± .0337, t(29.9)= 3.226, d = .770. **(F–G)** Scatter plots illustrating the relationship between RBCv and capillary diameter **(F)** and cell flux and capillary diameter **(G)**. Positive correlations in treated groups suggest improved capillary function. Plots of RBCv COV **(H)** and cell flux COV **(I). J)** Postprocessing of TPM angiograms using DeepVess. Each angiogram was composed of 100 slices, ∼50 µm below the top layer. Each stack was pre-processed for contrast enhancement (*left*) and then segmented into binary images and vessel center lines using DeepVess (*middle and right*)^43^ from where mean capillary segment diameter and standard deviation (SD) were estimated. Scale bar: 200 µm. Estimates were performed per mouse and per group. **K)** Plots for mean capillary segment diameter. **L)** Plots of mean capillary segment diameter SD. For comparisons a ranked LMM was used to account for non-normality. WT vs. Tg-SwDI, LMM, rank-diff = 22.8 ± 8.83, t(51.9)= 2.582, d = .950; Tg-SwDI vs. Tg-ATZ, LMM, rank-diff = –20.4 ± 8.67, t(52)= –2.355, d = .851; Tg-SwDI vs. Tg-MTZ, LMM, rank-diff = –18.9 ± 9.08, t(53)= –2.082, d = .788. **M)** Plot of mean capillary length density. Error bars = SD.

In WT mice, RBCv showed a positive correlation with capillary diameter (Fig. 3F), while cell flux did not (Fig. 3G). This indicates that autoregulation in healthy mice, likely mediated by pericytes^56^, ensures uniform red blood cell flux across capillaries, with velocity moderately scaling with diameter. In contrast, Tg-SwDI mice exhibited no correlation between RBCv or flux and capillary diameter, suggesting that such autoregulation is disrupted. The absence of correlation is hence indicative of morphological or pathological factors in Tg-SwDI mice that impede normal modulation of RBCv.

Finally, we calculated the coefficient of variance (COV) of RBCv and cell flux, reflecting capillary flow distribution (Fig. 3H and 3I). While Tg-SwDI mice exhibited a slightly higher cell flux COV compared to WT, this difference did not reach statistical significance, suggesting that capillary perfusion is maintained despite potential disturbances in capillary flows.

Collectively, these findings reveal early microvascular alterations in presymptomatic Tg- SwDI mice, including increased capillary stalling, reduced RBCv, and disrupted flow regulation. These changes, although subtle, suggest that early capillary dysfunction may precede overt reductions in blood flow or oxygen delivery, potentially contributing to the progression of AD-related pathology over time.

### 3.3 Disturbed capillary flow dynamics in presymptomatic Tg-SwDI are associated with alterations in capillary morphology

To examine whether capillary flow changes relate to altered capillary morphology, we examined TPM angiograms (Fig. 3J). The mean capillary diameter in Tg-SwDI mice showed no significant difference compared to WT mice (Fig. 3K and Supplementary Table 3). However, Tg-SwDI mice displayed increased diameter variability, as indicated by higher standard deviation (SD) (Fig. 3L) and coefficient of variance (COV) along capillary segments (Supplementary Fig. 4A and Supplementary Table 3).This variability suggests that Tg-SwDI mice may experience irregular capillary constrictions, possibly linked to pericyte activity as observed in animal models^57^ and ex-vivo studies^16^. These morphological disruptions may provide a structural basis for impaired capillary flow.

We also examined capillary tortuosity, which was similar in Tg-SwDI mice compared to WT mice (Supplementary Fig. 4B), aligning with observations in the APP/PS1 AD model^43^. Notably, sex-adjusted analysis revealed trends towards lower capillary length density (p = .0531, Cohen’s d = .818) (Fig. 3M), and lower capillary blood volume (p = .0783, Cohen’s d = .760, Supplementary Fig. 4C and Supplementary Table 3) in Tg-SwDI compared to WT mice. These finding suggest early structural changes that might contribute to flow abnormalities in the Tg-SwDI model.

### 3.4 Disturbed capillary blood flow dynamics and morphology are associated with increased CTH in presymptomatic Tg-SwDI mice

We measured mean transit-time (MTT) and CTH, the latter serving as an index of capillary dysfunction^58^, using indicator-dilution analysis developed for MRI in humans (Fig. 4A)^59^ and adopted for TPM (Fig. 4A-4C)^37^. MTT and CTH were measured between two parts of the microvasculature: one from a pial artery to a vein, representing the entire vascular network of the scanned region and best at capturing the vasculature detected within a human MRI voxel, and one from an arteriole to a venule, reflecting the hemodynamics of a smaller subset of capillaries (Fig. 4A).

**Figure 4.**
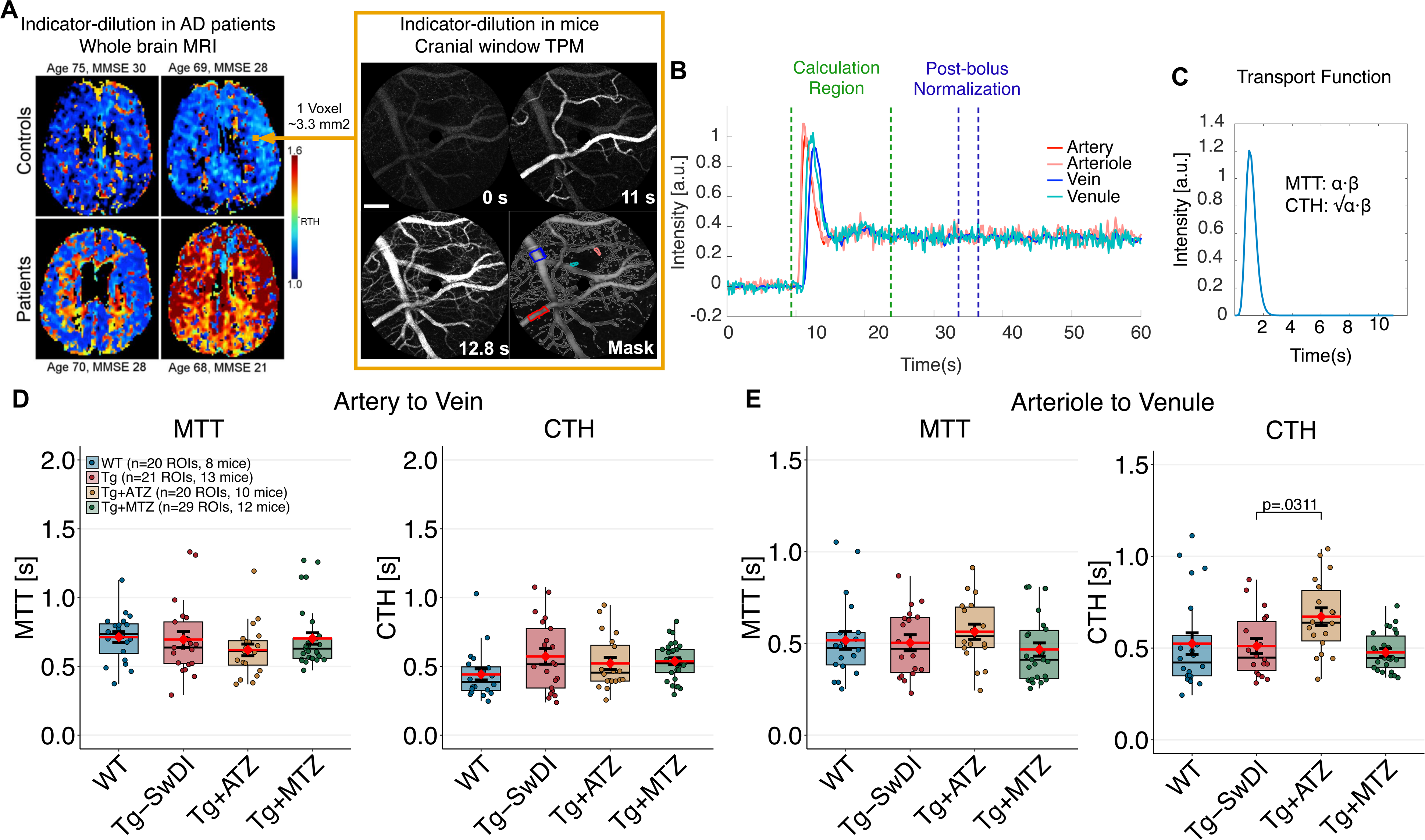
Capillary dysfunction is present in the Tg-SwDI but not in the treated mice with CAIs. **A).** Indicator-dilution method for estimating MTT and CTH in clinical settings using dynamic susceptibility contrast (DSC)-magnetic resonance imaging (MRI). Left: RTH maps (CTH/MTT) from two controls and two patients with different mini-mental scale evaluation (MMSE) scores predicting AD patients (Modified from Eskildsen et al.^92^) Maps of MTT and CTH are estimated in each voxel through parametric deconvolution^59^ between an arterial input function (AIF, i.e., middle cerebral artery) and a residue function (signal residue after the passage of the contrast agent). In contrast, high-spatial resolution of TPM (Right) allows the selection of AIF (red range colors) and a venous output function (VOF; blue range colors) within the field-of-view. Scans involved the injection of 10 µl of fluorescent dye while performing spiral scans (∼7 fps.). Scale bar: 100 µm. **B)** Intensity curves of vessels selected. A 10-s curve-matching region was selected to estimate MTT and CTH and all curves were normalized to the post-bolus baseline of the AIF. **C)** The probability transport function of capillary transit times h(t) was parameterized by a gamma variate with parameters α and β (MTT = α·β, CTH = √α·β). **D)** MTT and CTH estimations between arteries and veins. LMM were constructed to compare effect of treatment in MTT, CTH. **E)** MTT and CTH estimations between arterioles and venules. Tg-SwDI vs. Tg+ATZ log-diff. = .261 ± .103, t(82) = 2.530, d = .846. Error bars = SD.

A trend toward increased CTH was observed between arteries and veins in Tg-SwDI mice (LMM, p=.0572), with a large effect size (Cohen’s d=.9424), suggesting early-stage capillary dysfunction. A power test calculation estimated a sample size of 19 mice per group required to achieve statistical significance. While this trend fell short of conventional statistical significance, the effect size and alignment with elevated CTH in presymptomatic AD patients^19^, highlight its potential biological relevance. No differences in MTT were observed (Fig. 4E, Supplementary Table 4).

In passive compliant microvascular networks, MTT and CTH typically change in parallel as CBF increases, ensuring efficient oxygen extraction fraction, even at short capillary transit times^60^. However, at low CBF (long MTT), active homogenization of capillary flows is believed to cause a deviation from this relationship^61^, to ensure efficient oxygen extraction despite prolonged transit times. In our study, WT animals exhibited a low slope in the MTT-CTH relationship (Supplementary Fig. 5A), which we ascribe to this protective homogenization mechanism. In contrast, Tg-SwDI mice showed a lack of this compensatory mechanism, with higher CTH values at longer MTT, indicative of impaired microvascular flow regulation.

### 3.5 Disturbed capillary flow dynamics in presymptomatic Tg-SwDI mice do not affect oxygen extraction fraction (OEF)

To evaluate the impact of capillary flow disturbances on OEF, we measured in-vivo intravascular partial oxygen tension (pO_2_) using the oxygen-sensitive dye PtP-C343 with TPM phosphorescence lifetime (2PLM) as previously described^38,39^ (Fig. 5A-5B). Manual diameter estimations of scanned pial vessels at the region of 2PLM scans and segmentation of arteries, diving arterioles, upstream venules, and draining veins (Fig. 5C) revealed no discernible diameter differences between Tg-SwDI and WT (Fig. 5D and Supplementary Table 5).

**Figure 5.**
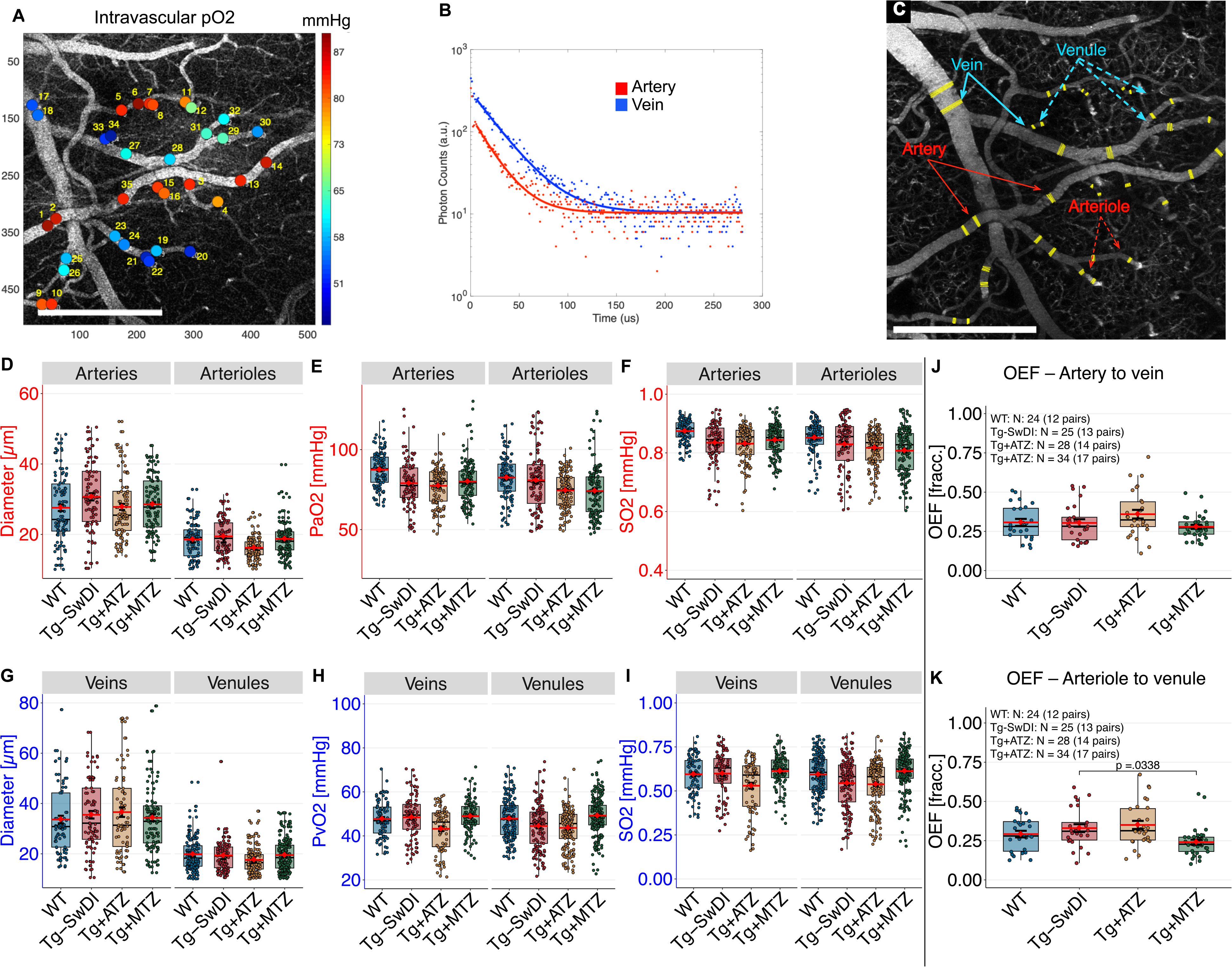
Alterations in OEF are not present in pre-symptomatic Tg-SwDI mice. **A)** intravascular (I.V.) pO_2_ was estimated using 2PLM by TPM after the I.V. injection of an oxygen-sensitive dye (PtP-C343). Single-point excitation was used, and photons were collected with a PMT and counted using time-correlated single-photon counting (TCSPC). Two single-point estimates were performed on each vessel from which MTT and CTH were estimated. **B)** PO_2_ was determined by fitting phosphorescence intensity decay with a single-exponential function using nonlinear least-squares method. Lifetime was converted to pO_2_ using a calibration plot from independent oxygen titration. Higher photon lifetime in arteries indicates higher O_2_ concentration compared to veins. **C)** Manual vessel diameter segmentation of vessels where pO_2_ was measured (Mean of 5 lines per point; N value per vessels type and group in Supplementary table 5). Estimates from arterial networks (arteries and diving arterioles): Diameter **(D)**, pO_2_ **(E)**, and SO_2_ **(F).** WT vs. Tg-SwDI Arterial SO_2_, LMM, rank-diff. = –152 ± 75.5, t(57.1) = –2.017, d = .818. Estimates from venous networks (veins and venules): Diameter **(G)**, pO_2_ **(I)**, and SO_2_ **(J)**. **K)** OEF between pial arteries and veins (mean per mouse, two scans per mouse). **L)** OEF between arterioles to venules. Tg-SwDI vs. Tg+MTZ, LMM, log-diff. = –.331 ± .134, t(57.2) = −2.461, d = 3.151. Error bars = SD.

We observed lower pO_2_ in the arteries and veins of male Tg-SwDI mice compared to females, and statistical models were thus adjusted by sex. Tg-SwDI mice exhibited trends towards lower pO_2_ (LMM, p=.0575, Fig. 5E) and oxygen saturation (SO_2_, LMM, p=.0570, Fig. 5F and Supplementary Table 6), supported by large effect sizes (pO_2_ Cohen’s d = .873, SO_2_ Cohen’s d=.872). A sample size of 22 mice per group is estimated to achieve statistical significance. These trends suggest potential systemic factors influencing cerebral oxygenation, such as compromised myocardial function, observed in AD patients^62^.

In WT mice, arterial pO_2_ increased with arterial diameter, explaining 15% of the variability in pO_2_ and SO_2_ across the arterial network (Supplementary Fig. 6A–B), consistent with previous studies in young WT (3-months-old)^63^. However, this correlation was absent in Tg-SwDI mice, implying other compensatory mechanisms, such as blood pressure regulation, may be influencing pO_2_ and SO_2_ levels in these mice. This observation aligns with reduced oxygen tension in brain parenchyma, a key regulator of oxygen diffusion^63^, previously reported in an AD mouse model^64^. No significant differences were found between WT and Tg-SwDI mice in the diameter, pO_2_, and SO_2_ of veins or venules (Fig. 5G – 5I).

Based on our pO_2_ measurements, we estimated OEF by pairing the mean PO_2_ of arteries and veins, and arterioles and venules, for each mouse. We observed higher OEF in males than females. After adjusting our analysis for sex effect, we did not observe differences in the averaged OEF per mouse between groups (Fig. 5J-5K).

### 3.6 Capillary stalling is associated with a higher expression of leukocyte adhesion molecules but not with BBB damage in presymptomatic Tg-SwDI mice

Previous studies have linked AD models and capillary stalling with increased expression of adhesion molecules like intracellular adhesion molecule 1 (ICAM-1)^15^, indicative of endothelial activation. Our results show elevated ICAM-1 levels in Tg-SwDI compared to WT, confirming endothelial activation (Fig. 6A). We also measured CypA molecular signature, linked to microvascular damage, including blood-brain barrier breakdown^65^, and assessed plasmatic levels of vascular endothelial growth factor A (VEGF-A), an essential pro-angiogenic factor associated with capillary stalling^66^. In contrast to the increase in ICAM-1, tissue CypA and plasmatic VEGF-A levels remained similar between groups (Fig. 6B and 6C).

**Figure 6.**
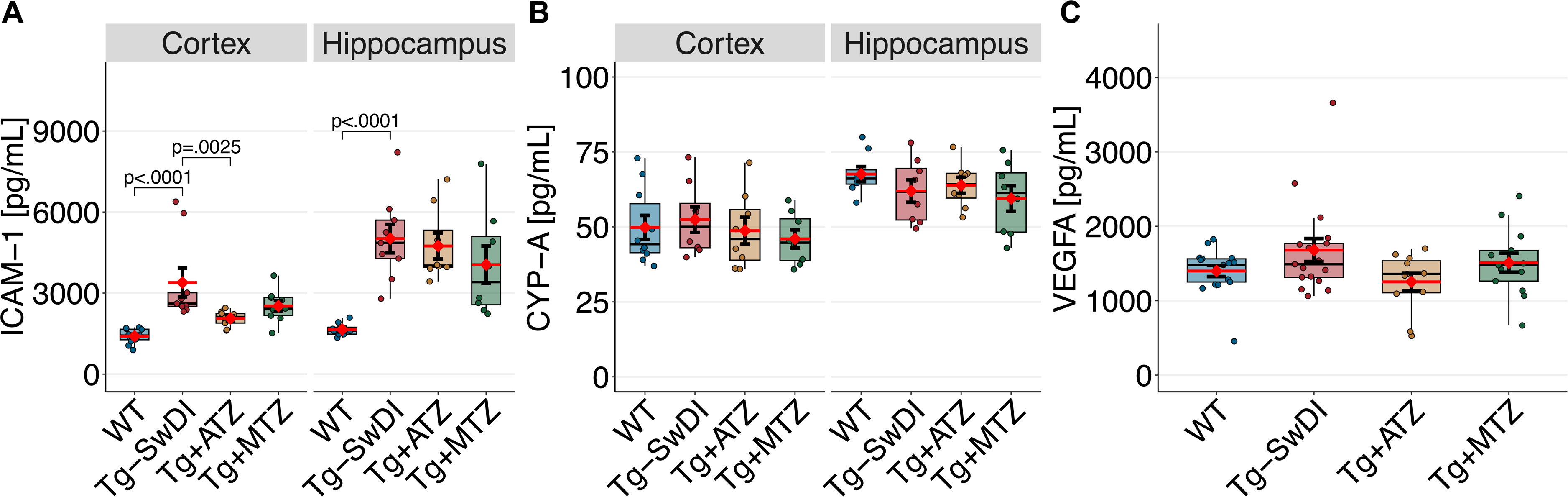
Leukocyte adhesion molecules and BBB marker levels. **A)** Box plot of ICAM- 1 levels in homogenized cortex and hippocampus, respectively, measured by ELISA. Significant difference between groups in ICAM-1 expression was observed in cortex (ANOVA, F[1, 35] = 10.046, p < .001) and hippocampus (F[1, 3] = 9.2224, p = .0019). Post-hoc pairwise comparisons using estimated marginal means for cortex: WT vs. Tg- SwDI log-diff. = .822 ± .114, t(35) = 7.206, d = 2.083; Tg-SwDI vs. Tg+ATZ log-diff. = – .429 ± .122, t(35) = 3.515, d = 1.086. Post-hoc for hippocampus: WT vs. Tg-SwDI diff. = 3369 ± 701 pg/ml, t(29) = 4.803, d = 1.727. **B)** Box plot of CYP-A levels obtained from ELISA analyses of homogenized cortex and hippocampus. N-value tissue ELISAs: WT: n=12 (5 F, 7 male); Tg-SwDI: n=9 mice (5 F, 4 male); Tg+ATZ: n=9 mice (5 F, 4 male; Tg+MTZ: n=9 mice (5 F, 4 male). **C)** Box plot of levels VEGF-A in plasma measured by Mesoscale. N-value tissue Mesoscale: WT: n=17 (7 F, 10 M); Tg-SwDI: n=9 mice (10 F, 7 M); Tg+ATZ: n=11 mice (7 F, 4 M; Tg+MTZ: n=13 mice (5 F, 8 M). Error bars = SD.

### 3.7 Effects of CAI Treatment on Cognitive Performance, Locomotion, and Brain Cortex Volume in Tg-SwDI Mice

We administered CAIs, ATZ (Tg+ATZ), and MTZ (Tg+MTZ) to separate groups of Tg- SwDI mice through the diet from 4 months old. Both CAIs were provided at dose of 20 mg/kg/day through the diet (100 ppm). Spatial reference memory, assessed with Barnes maze, showed no discernible effects of treatment at 9-10 months, noting that Tg-SwDI and WT animals showed similar memory functions at this time (Fig. 1A). Treatment with CAIs did not improved locomotor impairment compared to non-treated Tg-SwDI mice (Fig. 1B).

Notably, both treated groups did not show any difference in brain cortex volume compared to the Tg-SwDI mice. (Fig. 1D).

### 3.8 CAIs prevent capillary flow disturbances in presymptomatic Tg-SwDI mice

MTZ and ATZ show reduced cumulative duration of capillaries affected by stalling compared to untreated Tg-SwDI mice (Fig. 2H), with MTZ-treated mice also showing reduced stalling prevalence (Fig. 2G). Additionally, MTZ treatment led to significantly fewer prolonged capillary stalling events compared to Tg-SwDI controls (Kolmogorov-Smirnov test: p=.002, Supplementary Fig. 3C), suggesting a more localized and pronounced effect than ATZ in mitigating capillary stalling.

Both treatment groups showed enhanced single capillary hemodynamics, as evidence by increased RBCv and cell flux compared to untreated Tg-SwDI mice (Fig. 3B – 3C). Additionally, MTZ-treated mice showed larger capillary diameters (Fig. 3E), indicating enhanced capillary perfusion and reduced resistance. These findings suggest that CAI treatment alleviate capillary flow disturbances, potentially through improved capillary morphology or endothelial function.

Moreover, both treated groups show a proportional relationship between RBCv and capillary diameter, resembling the autoregulation observed in WT mice (Fig. 3F). This relationship suggest that CAIs may have a protective effect on capillary flow regulation, disrupted in non-treated mice. Enhanced cell flux in treated groups (Fig. 3G) underscores how increased capillary diameter improves perfusion, with MTZ-treated mice demonstrating a stronger effect. These findings underline the therapeutic potential of CAIs, particularly MTZ, in preserving capillary flow regulation of AD models.

### 3.9 CAIs prevent abnormalities in capillary morphology in presymptomatic Tg-SwDI mice

Neither MTZ nor ATZ treatment significantly altered capillary diameter (Fig. 3K). However, both reduced variability in diameter, as shown by decreased diameter SD (Fig. 3L) and COV (Supplementary Fig. 4A and Table 3). These reduced values suggest fewer narrowing regions along the capillary lumen, potentially reflecting improved capillary integrity.

The reduction in diameter variability in CAI-treated groups suggests fewer constricted capillary segments and improved pericyte regulation, though the modest changes raise questions about their attribution to treatment. Based on estimations from Nortley et al.^16^ (capillary diameter changes near pericytes) and Korte et al., (baseline diameters in AD), we calculated SDs of .182 µm (WT) and .207 µm (Tg-SwDI), reflecting a 13.73% relative increase in diameter variability in Tg-SwDI mice. Assuming pericytes influence 40 µm of their total 300 μm coverage^67^, variance was scaled proportionally. CAI treatment reduced variability modestly: 4.31% for Tg+ATZ (.051 µm) and 3.8% for Tg+MTZ (.045 µm), but changes remained modest compared to WT (Supplementary Methods 1). These results suggest that CAIs partially improve pericyte regulation but do not fully reverse amyloid-induced capillary dysfunction, likely due to limited pericyte influence and amyloid-driven capillary tone changes beyond direct regulation^16,57^.

Overall, these findings indicate that CAIs may help preserve capillary morphology in Tg- SwDI mice by reducing diameter variability along capillary segments. This reduction in variability could signify fewer constrictions within the capillary lumen, potentially linked to improved pericyte function.

### 3.10 MTZ treatment prevents capillary dysfunction and improves oxygen availability in presymptomatic Tg-SwDI mice

Treatment with CAIs showed differential effect on capillary flow distributions. While no significant differences were observed in MTT across groups (Fig. 4D-E and Supplementary Table 4), ATZ-treated mice exhibited a significant increase in CTH at arteriole-to-venule level (Fig. 4E), suggesting less stable capillary flow regulation in response to perfusion changes in the local capillary network. Conversely, MTZ-treated mice displayed reduced MTT-CTH correlation at the artery-to-vein, resembling the active flow regulation observed in WT mice (Supplementary Fig. 5). This active regulation supports consistent capillary transit times under variable flow conditions, contrasting the passive regulation observed in Tg-SwDI mice and partially in ATZ-treated mice (p=.0753).

In addition, preventing impaired capillary flow regulation, MTZ-treated mice showed lower OEF than untreated Tg-SwDI between arterioles and venules (Fig. 5J – 5K, and Supplementary Table 6). Despite reduction in capillary density (Fig. 3M), indictive of lower surface area for oxygen exchange, the increased RBCv and cell flux (Fig. 3C) and enlarged capillary diameters (Fig. 3E) in MTZ-treated mice likely enhance oxygen delivery efficiency. These changes, combined with shorter capillary stalls (Fig. 2H), suggest that MTZ treatment alleviated the negative effects of reduced capillary length density, ensuring adequate oxygen availability during steady-state conditions.

Altogether, these findings indicate that MTZ prevents capillary dysfunction and optimizes oxygen transport by enhancing flow dynamics, morphology, and delivery efficiency in presymptomatic Tg-SwDI mice.

### 3.11 CAIs reduce ICAM-1 expression and Aβ load in presymptomatic Tg-SwDI mice

ATZ-treated mice showed a significant reduction in ICAM-1 expression in cortex compared to non-treated Tg-SwDI mice, while MTZ-treated mice exhibited a trend toward reduced ICAM-1 expression (p=.0525, Cohen’s d=.6205), suggesting a weaker effect relative to ATZ (Fig. 6A). Both treatments exhibited no significant effects on CypA or VEGF-A expression (Fig. 6B–C)

In our previous study, CAI treatment prevented Aβ deposition in 15-month-old Tg-SwDI mice^27^. TPM measurements using Methoxy-X04 revealed a reduction of Aβ plaques in both treated groups. However, this reduction was statistically significant only in MTZ- treated mice, while ATZ-treated mice exhibited a trend toward reduction (p=.0717, Cohen’s d=.730; Fig. 1G). No significant changes were observed in total Aβ burden across groups (Fig. 1F). Post hoc analyses revealed a reduction in small plaques (Supplementary Fig. 7A) and a trend towards fewer very small (p=.0675, Tg+ATZ Cohen’s d=.155, Tg+MTZ Cohen’s d=.843, Supplementary Fig. 7A) and medium-sized plaques (p=.0677, Tg+ATZ Cohen’s d=.6895, Tg+MTZ Cohen’s d=.951, Supplementary Fig. 7C) in both groups. No difference in large plaques was observed (Supplementary Fig. 7D).

To further address the treatments’ impact on Aβ load, we quantified total Aβ-40 and Aβ-42 by conducting ELISA of homogenized cortex and hippocampus. Our analysis revealed lower levels of Aβ-40 and Aβ-42 load in the cortex and hippocampus of female mice. With sex-adjusted analysis, mice treated with ATZ exhibited a reduced Aβ-40 load in the cortex compared to non-treated mice and in the hippocampus relative to Tg-SwDI (Fig. 7A and Supplementary Table 7). MTZ-treated mice also showed tendency towards a lower Aβ-40 load in the cortex compared to Tg-SwDI (p=.0673), with a large effect (Cohen’s d=.811). Both treatment groups displayed a lower Aβ-42 load in the cortex (Fig. 7B and Supplementary Table 7). However, only the Tg+ATZ group showed a significant decreased in Aβ-42 load in the hippocampus (Fig. 7B).

**Figure 7.**
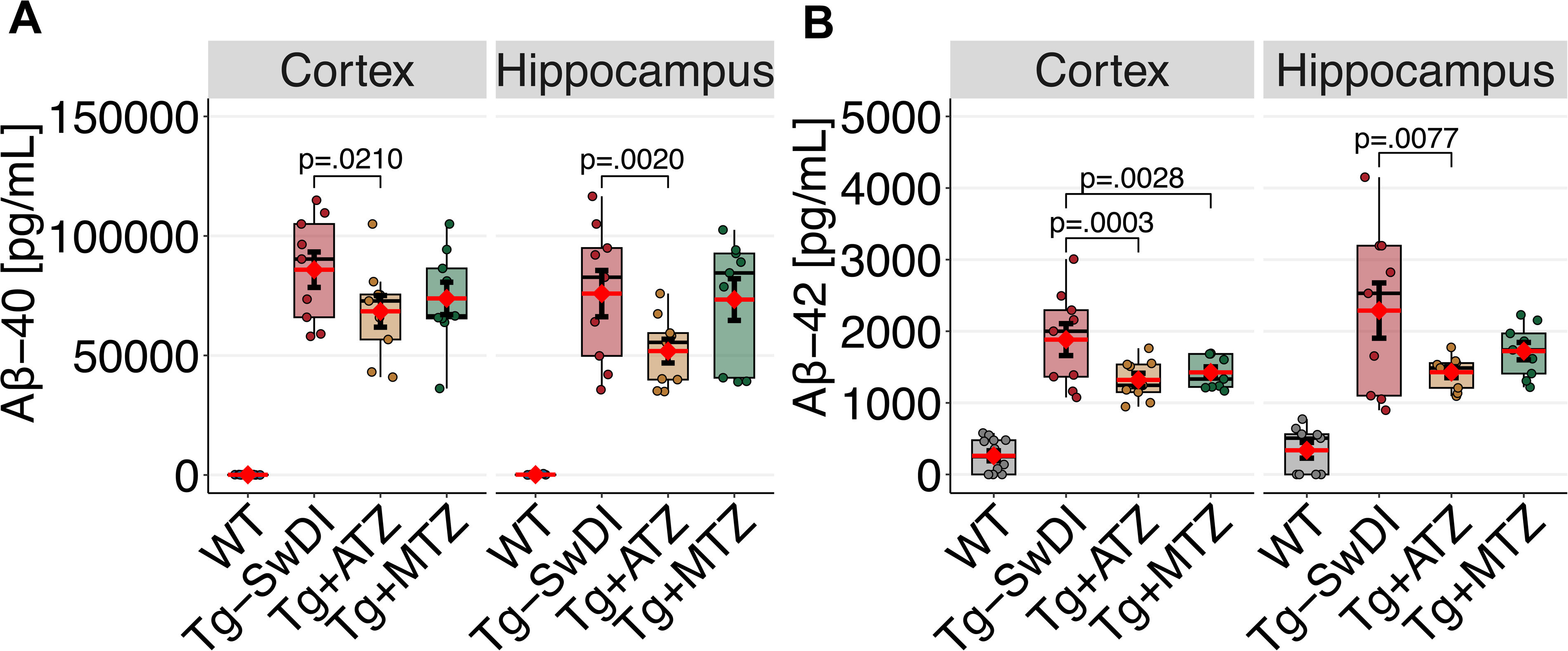
Amyloid overload is reduced in cortex and hippocampus in CAI treated Tg- SwDI mice. **A)** Box plot of total Aβ-40 load measured by ELISA of homogenized cortex and hippocampus, respectively. Significant difference between groups was observed in log-transformed Aβ-40 load in cortex (F[1, 2] = 4.0646) and hippocampus (F1, 2] = 3.14e9 pg/ml, p = .0011). Post-hoc pairwise comparisons using estimated marginal means for cortex: Tg-SwDI vs. Tg-ATZ diff. = –17416 ± 6253, t(23) = –2.785, d = 1.177. Pairwise comparisons for hippocampus: Tg-SwDI vs. Tg-ATZ log-diff. = –.343 ± .091, t(23) = – 3.762, d = 1.394. **B)** Box plot of total Aβ-42 load in cortex and hippocampus. Significant difference between groups was observed in log-transformed Aβ-42 load in cortex (F[1, 2] = 11.173, p < .001) and hippocampus (F[1, 2] = 6.9141 pg/ml, p = .0044). Post-hoc pairwise comparisons using estimated marginal means for cortex: Tg-SwDI vs. Tg+ATZ log-diff. = – .317 ± .069, t(23) = –4.567, d = 1.598; Tg-SwDI vs. Tg+MTZ log-diff. = –.232 ± .069, t(23) = –3.341, d 0 1.169. Pairwise comparison for hippocampus: Tg-SwDI vs. Tg+ATZ log-diff. = –.351 ± .109, t(23) = –3.214, d = 1.312. N-value for cortex and hippocampus: WT: n=12 (5 F, 7 male); Tg-SwDI: n=9 mice (5 F, 4 male); Tg+ATZ: n=9 mice (5 F, 4 male; Tg+MTZ: n=9 mice (5 F, 4 male). Error bars = SD.

## 4. Discussion

This study examined microvascular dynamics, capillary morphology, and Aβ load in presymptomatic, 9-10-month-old Tg-SwDI mice with cerebral amyloidosis. Our first finding reveals early microvascular changes, including capillary stalling, increased CTH, and capillary morphometric alterations, coexist with Aβ accumulation before cognitive impairment. These changes linked to endothelial activation, may reduce oxygen delivery as the model ages. Our second main finding is that long-term CAI treatment prevents microvascular disturbances, so treated mice had similar capillary morphometry to WT mice, reduced Aβ load, and fewer and shorter stalling events, possibly due to reduced endothelial activation.

Presymptomatic stages were defined by no memory impairment in Barnes maze testing. Discrepancies with prior studies in Tg-SwDI mice^68–72^ may result from methodological differences, such as maze design (8 vs. 20 holes maze) and outcome measures. Most deficits have only been observed in studies in which only trial latency has been reported. However, escape latency might be impacted by differences in factors independent of cognition such as decreases in motor function, motivation, anxiety or general search strategy^50,51^. In our study, escape latency did not differ between groups, suggesting that motor function did not affect performance. However, the Barnes maze alone may not fully capture subtle cognitive changes. Broader cognitive testing and larger cohorts are needed to confirm findings and evaluate vascular contributions to early AD stages in the Tg-SwDI model.

Studies have demonstrated that capillary flow disturbances appear in AD models during advanced disease stages^64,66,73^, and recent evidence shows that large vessel morphological changes and CBF reductions precede memory impairment in the 3xTg-AD model^74^. Our study extends these findings, showing that that capillary flow disturbances– marked by altered RBCv, cell flux, and capillary stalling–are present in presymptomatic mice with cerebral amyloidosis. This support the notion that capillary dysfunction may precede CBF reductions, similar to AD patients during the prodromal phases^18,75^. Longitudinal studies addressing capillary flow disturbances at stages of the disease are necessary to understand the progression of capillary flow disturbances during the development of AD.

Capillary flow disturbances in the Tg-SwDI mice are marked by increased CTH without changes in MTT, indicating impaired flow regulation despite preserved CBF. This dysfunction reflects disrupted oxygen extraction due to heterogenous capillary flow distribution^58^. In WT mice, active flow homogenization stabilizes CTH during hypoperfusion, ensuring consistent oxygen delivery across capillaries as previously predicted^14^. This mechanism, observed in prior studies with rats during hypovolemia^61^ and supported by our findings (Supplementary Fig. 5A), is absent in Tg-SwDI mice. Instead, Tg-SwDI mice exhibit passive flow regulation, as shown by the linear relationship between CTH and MTT^60^. This lack of active adjustments heightens the brain’s vulnerability to oxygen deficits during CBF reductions. These findings highlight the need to study CTH and MTT together to understand capillary dysfunction in AD.

Capillary dysfunction in AD likely involves multiple factors, including hematocrit, capillary diameter, length, and vessel stiffness^76^. In WT and treated groups, RBCv and cell flux correlate with capillary diameter, highlighting the role of diameter in maintaining effective perfusion^77^. However, this correlation is absent in Tg-SwDI mice, suggesting structural or functional alterations in capillary walls. Evidence of pericyte constriction^16^ and dysfunction^78^ in AD supports the hypothesis that flow disturbances result from a combination of capillary stalls and pericyte rigidity or impairment.

Morphometric data show increased variability in capillary diameter in Tg-SwDI mice, indicating narrowed segments that may disrupt flow and increase resistance (Fig. 8). These narrowed regions restrict leukocyte passage, forcing them to deform into disk-like shapes and increasing their contact to the capillary wall as previously modelled^79^. This elevated contact surface likely promotes the expression of adhesion molecules like ICAM- 1, as observed in our study. Increased ICAM-1 expression exacerbates leukocyte adhesion, compounding resistance and further impairing capillary perfusion. Figure 8 illustrates how these structural changes contribute to capillary dysfunction by diverting blood flow and reducing perfusion efficiency. Collectively, these findings highlight capillary narrowing as a key driver of flow disturbances in presymptomatic Tg-SwDI mice.

**Figure 8.**
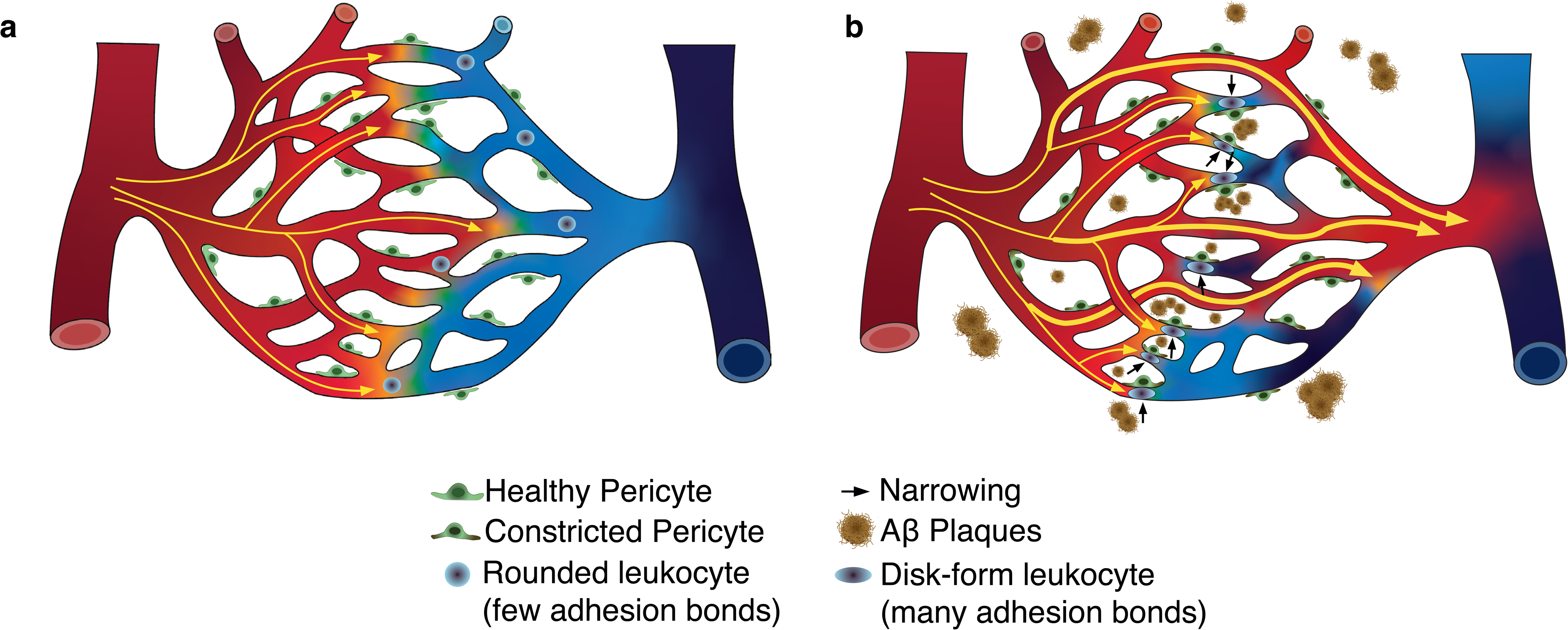
Disturbances in capillary patterns in Tg-SwDI induce capillary dysfunction. **A)** In healthy capillary networks, the transport of oxygen between blood and tissue is limited by blood’s capillary transit time and the distribution of capillary flows (thin yellow arrows) at a given time. Leukocytes commonly have a rounded shape with few adhesion bonds not affecting homogenization of capillary flows. **B)** Phenomena affecting capillary patency, such as lumen narrowing and stalling, can disturb capillary flow patterns. In AD narrowing is suggested to occur at regions at the pericyte soma^16^. Consequently, leukocytes will need to squeeze through the capillaries, taking on a disk shape and increasing the number of adhesion bonds, which promotes capillary stalling. These pathological phenomena induce shunting of capillary flows through low-resistance capillaries (thick yellow paths/arrows) and prevent their normal homogenization (increased CTH), leading to capillary dysfunction during increases in blood flow.

Reductions in pO_2_ and SO_2_ in pial arteries suggest factors influencing cerebral oxygenation, from local to systemic origin. Locally, this may involve the contribution of large vessels, which account for ∼50% of extracted O_2_ during steady-state^63^ Tissue pO₂ measurements in AD mouse models have demonstrated a reduction in tissue pO₂ compared to WT mice, accompanied by an increase in OEF^80^. The elevated OEF reflects a compensatory response to impaired capillary flow regulation, aiming to preserve oxygen delivery despite vascular dysfunction. However, this compensatory mechanism can lead to decreased arterial SO₂ levels within the microvasculature and tissue, ultimately reflecting impaired oxygenation in AD brains.

Systemic factors, such as cardiac function or gas exchange impairments in the lungs, may also contribute to these changes in intravascular oxygen levels. Cardiac dysfunction, including Aβ deposition in the myocardium, has been previously reported in AD patients^62^ and the Tg2576 AD mouse model^81^, underscoring the role of brain-heart interactions in AD. Additionally, respiratory impairment has been linked to damage in the brainstem, where nuclei involved in breath control are located^82,83^. While evidence of disturbances in basal ventilation in AD animal models is scarce, respiratory impairment has been observed in the Tau-P301L mouse model of tauopathy^84^. Although these changes may not directly cause mortality, they can contribute to hypoxia and reduce SO₂ levels locally. Together, these findings emphasize the synergistic impact of systemic and local impairments, where cardiac and capillary dysfunction amplify each other’s effects on brain perfusion. Further studies assessing tissue and intravascular oxygen levels, along with evaluations of cardiac and respiratory function, are needed to fully understand the mechanisms underlying reductions in pO₂ and SO₂ at the pial vasculature.

We demonstrate that CAIs prevent the development capillary flow disturbances, as evidenced by shorter capillary stalls, increased RBCv and cell flux, and reduced variability in capillary diameter in treated Tg-SwDI mice, with no difference in diameter of pial vasculature. MTZ treatment further showed larger capillary diameters, reduced OEF, and decreased capillary density, suggesting improved oxygen delivery that may limit hypoxia-driven angiogenesis. The lower OEF in MTZ-tread mice likely reflects increased CBF and reduced CTH, enhancing oxygen delivery efficiency, as predicted by Jespersen & Østergaard^14^ and supported by indicator-dilution methods^37^. In healthy vascular networks, like in WT mice, reductions in CTH compensate for increased MTT, maintaining efficient oxygen extraction despite reduced CBF^14^. MTZ-treated mice showed similar adjustments of CTH whenever MTT increases, unlike non-treated Tg-SwDI and ATZ-treated mice, indicating minimal active regulation. These findings suggest that MTZ preserved capillary flow regulation and oxygen availability, mitigating the effect of capillary dysfunction in Tg- SwDI.

Previous studies have examined vasodilator treatments in AD models, such as nicorandil, which targets nitrate content and potassium channel activation, but these showed no improvement in CBF in the 3xTg AD model^74^. Unlike nicorandil, CAIs demonstrate benefits that extend beyond vasodilation. They prevent mitochondrial dysfunction, inhibit pro-apoptotic pathways, reduce glial reactivity markers, and activate anti-inflammatory microglia^20,26,27^, creating a healthier brain environment that promotes efficient Aβ degradation. These effects aligns with the reduced Aβ load observed in this and prior research^27^. Further, evidence suggests MTZ directly activates angiotensin-converting enzyme II (ACE2), producing angiotensin-(1-7), a potent vasodilator associated with Mas receptor activation^85^. This mechanism may enhance vascular tone and perfusion, contributing to the protective effects of MTZ. CAIs may also directly influence pericyte survival; inhibitions of mitochondrial carbonic anhydrases (CA-VA and -VB) has been shown to prevent glucose-mediated pericyte apoptosis in diabetic models^86,87^. Elevated CA-VB levels in the Tg-SwDI^27^, suggest that CAIs could counteract amyloid-driven capillary dysfunction by supporting pericyte health. With our current animal model, we could not visualize pericytes, which prevented us from addressing their precise role in these mechanisms. Employing transgenic AD models with fluorescently labelled pericytes, as used in recent studies^78^, could allow precise examination of pericyte-mediated diameter changes and further evaluate their contribution to capillary dysfunction and the therapeutic efficacy of CAIs. Even without this information, our findings position CAIs as promising therapeutic candidates for mitigating capillary flow disturbances and broader cerebrovascular pathologies in AD.

Notably, we observed that treatment with ATZ showed an ambivalent effect, preventing Aβ load but with limited improvement in capillary hemodynamics. This discrepancy may be attributed to known secondary effects of ATZ, such as its association with impaired diaphragmatic muscle function and reduced exercise capacity^88^, and an increased risk of renal failure^89^. Such adverse health effects might negatively impact vascular integrity and functional outcomes. MTZ, on the other hand, demonstrated a more robust impact on both capillary hemodynamics and Aβ pathology, potentially due to its additional mechanisms, such as ACE2 activation, which may amplify its vasoprotective effects. The divergence in the efficacy of these two CAIs highlights the importance of considering individual drug properties and off-target effects in the development of therapeutic strategies for AD.

### Study limitations

The temporal resolution of OCTA imaging (∼8.5 seconds per volume) may miss brief capillary stalls under one frame, potentially underestimating transient events. Conversely, including all stalls regardless of duration could inflate the representation of very short no-flow events. Despite these limitations, our uniform and unbiased analysis across experimental groups ensured valid comparisons. Future studies integrating higher temporal resolution techniques or complementary flow measurements could further refine these observations.

Other limitations include a modest sample size and the absence of tau pathology. Future research with larger cohorts and tau-inclusive models is needed to enhance findings.

Although systemic CAIs like ATZ and MTZ are widely used, side effects limit long-term use^23^, warranting development of selective CA inhibitors and optimized dosing. To address these limitations, developing more selective CA inhibitors targeting specific enzymes or mechanisms, and optimizing dosing regimens to minimize side effects, may be necessary strategies for clinical applications^20,90,91^, see also U.S. Patent No. 10780094B2.

### Conclusion

Our findings highlight capillary flow disturbances and dysfunction as an early biomarker and therapeutic target in AD. Long-term CAI treatment, particularly MTZ, prevents capillary dysfunction, brain remodelling, and amyloid load, positioning CAIs as promising treatments to slow AD progression.

## Authors contributions

EGJ, SF, and LØ conceived the study. EGJ established the study objectives, designed and performed the experiments, acquired, and processed TPM data, statistical analysis, and wrote the first draft of the manuscript. PMR, IKM, and SS set up and optimized TPM imaging processing software, supported in establishing TPM sequences and interpretation of data analysis. Further, PMR supported in statistical analysis. SK and DB supported in OCT acquisition software and imaging post-processing. SF supported in training of mice and scanning sessions. BH performed, analyzed, and interpreted MRI imaging. LB established awake imaging in the laboratory and monitoring software together with EGJ and SF. MEK and SV synthesized and calibrated the oxygen probe PtP-C343. JC and JP performed molecular biology tests, data analysis and interpretation. JRC reviewed data analysis, the manuscript and supported in interpretation of data. BW and DLG performed analysis of OCT data. EC, ACM, and BGL performed behavior tests and processing of behavior data. SF contributed to study objectives, experimental design, interpretation of data and critically reviewed the manuscript. LØ mentored EGJ and contributed to the study design, objectives, and interpretation of data and critically evaluated the manuscript.

## Acknowledgements

We would like to thank you Nina K. Iversen for supporting on the randomization of the treatment. We would like to thank Morten Frank Andersen for his support in segmenting the MRI data and Claudia Martinez for segmenting pial vessels from TPM data. We also thank Susanne S. Christensen and Anna Bay for their technical support and administration of the laboratory resources.

## Data and Software Availability

Data for each plot and associated scripts: https://zenodo.org/uploads/14285647. Macros for preprocessing angiograms and stacks for amyloid quantification, and DeepVess postprocessing scripts: https://zenodo.org/records/13762334. Segmentation model for amyloid plaques: https://zenodo.org/records/14285562. Software utilized in prior studies is available upon request. If additional information or clarification regarding any tool or method is needed, we would be pleased to provide it.

## Consent Statement

Human consent was not necessary for the current study.

## References

1. Alzheimer’s Disease International (ADI), Wimo A, Prince M, International AD. World Alzheimer Report 2015, The Global Impact of Dementia. Alzheimer’s Disease International (ADI). Published online 2015. doi:10.1111/j.0963-7214.2004.00293.x

2. Europe A. Dementia in Europe Yearbook 2019 Estimating the prevalence of dementia in Europe. Published online 2020.

3. Jack CR, Knopman DS, Jagust WJ, et al. Tracking pathophysiological processes in Alzheimer’s disease: an updated hypothetical model of dynamic biomarkers. The Lancet Neurology. 2013;12(2):207–216. doi:10.1016/S1474-4422(12)70291-0

4. Aisen PS, Cummings J, Jack CR, et al. On the path to 2025: understanding the Alzheimer’s disease continuum. Alz Res Therapy. 2017;9(1):60. doi:10.1186/s13195-017-0283-5

5. Bateman RJ, Smith J, Donohue MC, et al. Two Phase 3 Trials of Gantenerumab in Early Alzheimer’s Disease. N Engl J Med. 2023;389(20):1862–1876. doi:10.1056/NEJMoa2304430

6. Van Dyck CH, Swanson CJ, Aisen P, et al. Lecanemab in Early Alzheimer’s Disease. N Engl J Med. 2023;388(1):9–21. doi:10.1056/NEJMoa2212948

7. Rabin JS, Nichols E, La Joie R, et al. Cerebral amyloid angiopathy interacts with neuritic amyloid plaques to promote tau and cognitive decline. Brain. 2022;145(8):2823–2833. doi:10.1093/brain/awac178

8. Iturria-Medina Y, Sotero RC, Toussaint PJ, et al. Early role of vascular dysregulation on late-onset Alzheimer’s disease based on multifactorial data-driven analysis. Nature Communications. 2016;7(1):1–14. doi:10.1038/ncomms11934

9. Iadecola C. The overlap between neurodegenerative and vascular factors in the pathogenesis of dementia. Acta Neuropathologica. 2010;120(3):287–296.

10. Carey A, Fossati S. Hypertension and hyperhomocysteinemia as modifiable risk factors for Alzheimer’s disease and dementia: New evidence, potential therapeutic strategies, and biomarkers. Alzheimer’s & Dementia. 2023;19(2):671–695. doi:10.1002/alz.12871

11. Gutiérrez-Jiménez E, Angleys H, Rasmussen PM, et al. Disturbances in the control of capillary flow in an aged APPswe/PS1ΔE9 model of Alzheimer’s disease. Neurobiology of Aging. 2018;62:82–94. doi:10.1016/j.neurobiolaging.2017.10.006

12. Nation DA, Sweeney MD, Montagne A, et al. Blood-brain barrier breakdown is an early biomarker of human cognitive dysfunction. Nature Medicine. 2019;25:270–276. doi:10.1038/s41591-018-0297-y

13. Iadecola C. The Pathobiology of Vascular Dementia. Neuron. 2013;80(4):844–866. doi:10.1016/j.neuron.2013.10.008

14. Jespersen SN, Østergaard L. The roles of cerebral blood flow, capillary transit time heterogeneity, and oxygen tension in brain oxygenation and metabolism. Journal of Cerebral Blood Flow and Metabolism. 2012;32(2):264–277. doi:10.1038/jcbfm.2011.153

15. Cruz Hernández JC, Bracko O, Kersbergen CJ, et al. Neutrophil adhesion in brain capillaries reduces cortical blood flow and impairs memory function in Alzheimer’s disease mouse models. Nature Neuroscience. 2019;22(3):413–420. doi:10.1038/s41593-018-0329-4

16. Nortley R, Korte N, Izquierdo P, et al. Amyloid b oligomers constrict human capillaries in Alzheimer’s disease via signaling to pericytes. Science. 2019;365(6450). doi:10.1126/science.aav9518

17. Østergaard L, Aamand R, Gutiérrez-Jiménez E, et al. The capillary dysfunction hypothesis of Alzheimer’s disease. Neurobiology of Aging. 2013;34(4):1018–1031. doi:10.1016/j.neurobiolaging.2012.09.011

18. Nielsen RB, Parbo P, Ismail R, et al. Impaired perfusion and capillary dysfunction in prodromal Alzheimer’s disease. Alzheimer’s &amp; Dementia: Diagnosis, Assessment &amp; Disease Monitoring. 2020;12(1). doi:10.1002/dad2.12032

19. Madsen LS, Kjeldsen PL, Ismail R, et al. Capillary dysfunction in healthy elderly *APOE ε* 4 carriers with raised brain Aβ deposition. Alzheimer’s &amp; Dementia. 2024;20(1):459–471. doi:10.1002/alz.13461

20. Lemon N, Canepa E, Ilies MA, Fossati S. Carbonic Anhydrases as Potential Targets Against Neurovascular Unit Dysfunction in Alzheimer’s Disease and Stroke. Frontiers in Aging Neuroscience. 2021;13(November):1–19. doi:10.3389/fnagi.2021.772278

21. Provensi G, Carta F, Nocentini A, et al. A new kid on the block? Carbonic anhydrases as possible new targets in alzheimer’s disease. International Journal of Molecular Sciences. 2019;20(19). doi:10.3390/ijms20194724

22. Supuran CT, Altamimi ASA, Carta F. Carbonic anhydrase inhibition and the management of glaucoma: a literature and patent review 2013-2019. Expert Opinion on Therapeutic Patents. 2019;29(10):781–792. doi:10.1080/13543776.2019.1679117

23. Wright A, Brearey S, Imray C. High hopes at high altitudes: pharmacotherapy for acute mountain sickness and high-altitude cerebral and pulmonary oedema. Expert Opinion on Pharmacotherapy. 2008;9(1):119–127. doi:10.1517/14656566.9.1.119

24. Supuran CT. Carbonic anhydrases: novel therapeutic applications for inhibitors and activators. Nat Rev Drug Discov. 2008;7(2):168–181. doi:10.1038/nrd2467

25. Jensen FB. Red blood cell pH, the Bohr effect, and other oxygenation-linked phenomena in blood O_2_ and CO_2_ transport. Acta Physiologica Scandinavica. 2004;182(3):215–227. doi:10.1111/j.1365-201X.2004.01361.x

26. Anzovino A, Canepa E, Alves M, Lemon NL, Carare RO, Fossati S. Amyloid Beta Oligomers Activate Death Receptors and Mitochondria-Mediated Apoptotic Pathways in Cerebral Vascular Smooth Muscle Cells; Protective Effects of Carbonic Anhydrase Inhibitors. Cells. 2023;12(24):2840. doi:10.3390/cells12242840

27. Canepa E, Parodi-Rullan R, Vazquez-Torres R, et al. FDA-approved carbonic anhydrase inhibitors reduce amyloid β pathology and improve cognition, by ameliorating cerebrovascular health and glial fitness. Alzheimer’s & Dementia. Published online April 26, 2023:alz.13063. doi:10.1002/alz.13063

28. Setti SE, Flanigan T, Hanig J, Sarkar S. Assessment of sex-related neuropathology and cognitive deficits in the Tg-SwDI mouse model of Alzheimer’s disease. Behavioural Brain Research. 2022;428:113882. doi:10.1016/j.bbr.2022.113882

29. Park L, Koizumi K, El Jamal S, et al. Age-dependent neurovascular dysfunction and damage in a mouse model of cerebral amyloid angiopathy. Stroke; a journal of cerebral circulation. 2014;45(6):1815–1821.

30. Bach ME, Hawkins RD, Osman M, Kandel ER, Mayford M. Impairment of Spatial but Not Contextual Memory in CaMKII Mutant Mice with a Selective Loss of Hippocampal LTP in the Range of the 0 Frequency.

31. Pitts MW. Barnes Maze Procedure for Spatial Learning and Memory in Mice. Bio Protoc. 2018;8(5):e2744. doi:10.21769/BioProtoc.2744

32. Desjardins M, Kılıç K, Thunemann M, et al. Awake Mouse Imaging: From Two-Photon Microscopy to Blood Oxygen Level–Dependent Functional Magnetic Resonance Imaging. Biological Psychiatry: Cognitive Neuroscience and Neuroimaging. 2019;4(6):533–542. doi:10.1016/j.bpsc.2018.12.002

33. Lindhardt TB, Gutiérrez-Jiménez E, Liang Z, Hansen B. Male and Female C57BL/6 Mice Respond Differently to Awake Magnetic Resonance Imaging Habituation. Frontiers in Neuroscience. 2022;16. Accessed January 11, 2023. https://www.frontiersin.org/articles/10.3389/fnins.2022.853527

34. Fruekilde SK, Bailey CJ, Lambertsen KL, et al. Disturbed microcirculation and hyperaemic response in a murine model of systemic inflammation. J Cereb Blood Flow Metab. 2022;42(12):2303–2317. doi:10.1177/0271678X221112278

35. Srinivasan VJ, Jiang JY, Yaseen MA, et al. Rapid volumetric angiography of cortical microvasculature with optical coherence tomography. Optics Letters. 2010;35(1):43. doi:10.1364/OL.35.000043

36. Erdener ŞE, Tang J, Sajjadi A, et al. Spatio-temporal dynamics of cerebral capillary segments with stalling red blood cells. Journal of Cerebral Blood Flow and Metabolism. 2019;39(5):886–900. doi:10.1177/0271678X17743877

37. Gutiérrez-Jiménez E, Cai C, Mikkelsen IK, et al. Effect of electrical forepaw stimulation on capillary transit-time heterogeneity (CTH). Journal of Cerebral Blood Flow and Metabolism. Published online 2016. doi:10.1177/0271678X16631560

38. Finikova OS, Lebedev AY, Aprelev A, et al. Oxygen Microscopy by Two-Photon-Excited Phosphorescence. Chemphyschem. 2008;9(12):1673–1679. doi:10.1002/cphc.200800296

39. Sakadžić S, Roussakis E, Yaseen MA, et al. Two-photon high-resolution measurement of partial pressure of oxygen in cerebral vasculature and tissue. Nature Methods. Published online 2010. doi:10.1038/nmeth.1490

40. Li B, Esipova TV, Sencan I, et al. More homogeneous capillary flow and oxygenation in deeper cortical layers correlate with increased oxygen extraction. eLife. Published online 2019. doi:10.7554/eLife.42299

41. Drew PJ, Blinder P, Cauwenberghs G, Shih AY, Kleinfeld D. Rapid determination of particle velocity from space-time images using the Radon transform. J Comput Neurosci. 2010;29(1- 2):5–11. doi:10.1007/s10827-009-0159-1

42. Kleinfeld D, Mitra PP, Helmchen F, Denk W. Fluctuations and stimulus-induced changes in blood flow observed in individual capillaries in layers 2 through 4 of rat neocortex. Proc Natl Acad Sci USA. 1998;95(26):15741–15746. doi:10.1073/pnas.95.26.15741

43. Haft-Javaherian M, Fang L, Muse V, Schaffer CB, Nishimura N, Sabuncu MR. Deep convolutional neural networks for segmenting 3D in vivo multiphoton images of vasculature in Alzheimer disease mouse models. PLOS ONE. 2019;14(3):e0213539. doi:10.1371/JOURNAL.PONE.0213539

44. Berg S, Kutra D, Kroeger T, et al. ilastik: interactive machine learning for (bio)image analysis. Nature Methods 2019 16:12. 2019;16(12):1226-1232. doi:10.1038/s41592-019-0582-9

45. Ardalan M, Chumak T, Quist A, et al. Reelin cells and sex-dependent synaptopathology in autism following postnatal immune activation. British J Pharmacology. 2022;179(17):4400–4422. doi:10.1111/bph.15859

46. Yushkevich PA, Piven J, Hazlett HC, et al. User-guided 3D active contour segmentation of anatomical structures: Significantly improved efficiency and reliability. NeuroImage. 2006;31(3):1116–1128. doi:10.1016/j.neuroimage.2006.01.015

47. Mowbray FI, Fox-Wasylyshyn SM, El-Masri MM. Univariate Outliers: A Conceptual Overview for the Nurse Researcher. Can J Nurs Res. 2019;51(1):31–37. doi:10.1177/0844562118786647

48. Bates D, Mächler M, Bolker BM, Walker SC. Fitting Linear Mixed-Effects Models Using lme4. Journal of Statistical Software. 2015;67(1):1–48. doi:10.18637/JSS.V067.I01

49. Sullivan GM, Feinn R. Using Effect Size—or Why the *P* Value Is Not Enough. Journal of Graduate Medical Education. 2012;4(3):279–282. doi:10.4300/JGME-D-12-00156.1

50. Gawel K, Gibula E, Marszalek-Grabska M, Filarowska J, Kotlinska JH. Assessment of spatial learning and memory in the Barnes maze task in rodents—methodological consideration. Naunyn-Schmiedeberg’s Arch Pharmacol. 2019;392(1):1–18. doi:10.1007/s00210-018-1589-y

51. O’Leary TP, Brown RE. Optimization of apparatus design and behavioral measures for the assessment of visuo-spatial learning and memory of mice on the Barnes maze. Learn Mem. 2013;20(2):85–96. doi:10.1101/lm.028076.112

52. Li M, Kitamura A, Beverley J, et al. Impaired Glymphatic Function and Pulsation Alterations in a Mouse Model of Vascular Cognitive Impairment. Front Aging Neurosci. 2022;13:788519. doi:10.3389/fnagi.2021.788519

53. Hébert F, Grand’Maison M, Ho MK, Lerch JP. Cortical atrophy and hypoperfusion in a transgenic mouse model of Alzheimer’s disease. Neurobiology of łdots. Published online 2013.

54. Phan TX, Baratono S, Drew W, et al. Increased Cortical Thickness in Alzheimer’s Disease. Annals of Neurology. 2024;95(5):929–940. doi:10.1002/ana.26894

55. Munting LP, Derieppe M, Suidgeest E, et al. Cerebral blood flow and cerebrovascular reactivity are preserved in a mouse model of cerebral microvascular amyloidosis. eLife. 2021;10:e61279. doi:10.7554/eLife.61279

56. Hall CN, Reynell C, Gesslein B, et al. Capillary pericytes regulate cerebral blood flow in health and disease. Nature. Published online 2014. doi:10.1038/nature13165

57. Korte N, Nortley R, Attwell D. Cerebral blood flow decrease as an early pathological mechanism in Alzheimer’s disease. Acta Neuropathologica. 2020;140(6):793–810. doi:10.1007/s00401-020-02215-w

58. Østergaard L, Jespersen SN, Engedahl T, et al. Capillary Dysfunction: Its Detection and Causative Role in Dementias and Stroke. Current Neurology and Neuroscience Reports. 2015;15(6):37. doi:10.1007/s11910-015-0557-x

59. Mouridsen K, Hansen MB, Østergaard L, Jespersen SN. Reliable estimation of capillary transit time distributions using DSC-MRI. Journal of Cerebral Blood Flow and Metabolism. 2014;34(9):1511–1521. doi:10.1038/jcbfm.2014.111

60. Rasmussen PM, Jespersen SN, Østergaard L. The effects of transit time heterogeneity on brain oxygenation during rest and functional activation. Journal of Cerebral Blood Flow and Metabolism. Published online 2015. doi:10.1038/jcbfm.2014.213

61. Hudetz AG, Feher G, Weigle CG, Knuese DE, Kampine JP. Video microscopy of cerebrocortical capillary flow: response to hypotension and intracranial hypertension. American Journal of Physiology-Heart and Circulatory Physiology. 1995;268(6):H2202–H2210. doi:10.1152/ajpheart.1995.268.6.H2202

62. Troncone L, Luciani M, Coggins M, et al. Aβ Amyloid Pathology Affects the Hearts of Patients With Alzheimer’s Disease. Journal of the American College of Cardiology. 2016;68(22):2395–2407. doi:10.1016/j.jacc.2016.08.073

63. Sakadžić S, Mandeville ET, Gagnon L, et al. Large arteriolar component of oxygen delivery implies a safe margin of oxygen supply to cerebral tissue. Nature Communications. 2014;5. doi:10.1038/ncomms6734

64. Lu X, Moeini M, Li B, et al. A Pilot Study Investigating Changes in Capillary Hemodynamics and Its Modulation by Exercise in the APP-PS1 Alzheimer Mouse Model. Frontiers in Neuroscience. 2019;13. Accessed March 6, 2024. https://www.frontiersin.org/journals/neuroscience/articles/10.3389/fnins.2019.01261

65. Bell RD, Winkler EA, Singh I, et al. Apolipoprotein e controls cerebrovascular integrity via cyclophilin A. Nature. 2012;485(7399):512-516. doi:10.1038/nature11087

66. Ali M, Falkenhain K, Njiru BN, et al. VEGF signalling causes stalls in brain capillaries and reduces cerebral blood flow in Alzheimer’s mice. Brain. 2022;145(4):1449–1463. doi:10.1093/brain/awab387

67. Hartmann DA, Underly RG, Grant RI, Watson AN, Lindner V, Shih AY. Pericyte structure and distribution in the cerebral cortex revealed by high-resolution imaging of transgenic mice. Neurophoton. 2015;2(4):041402. doi:10.1117/1.NPh.2.4.041402

68. Xu F, Grande AM, Robinson JK, et al. Early-onset subicular microvascular amyloid and neuroinflammation correlate with behavioral deficits in vasculotropic mutant amyloid β-protein precursor transgenic mice. Neuroscience. Published online 2007. doi:10.1016/j.neuroscience.2007.01.043

69. Fan R, DeFilippis K, Van Nostrand WE. Induction of complement proteins in a mouse model for cerebral microvascular Aβ deposition. J Neuroinflammation. 2007;4(1):22. doi:10.1186/1742-2094-4-22

70. Xu F, Kotarba AME, Ou-Yang MH, et al. Early-onset formation of parenchymal plaque amyloid abrogates cerebral microvascular amyloid accumulation in transgenic mice. Journal of Biological Chemistry. 2014;289(25):17895–17908.

71. Anderson M, Xu F, Ou-Yang MH, Davis J, Van Nostrand WE, Robinson JK. Intensive ‘Brain Training’ Intervention Fails to Reduce Amyloid Pathologies or Cognitive Deficits in Transgenic Mouse Models of Alzheimer’s Disease. JAD. 2017;55(3):1109–1121. doi:10.3233/JAD-160674

72. Robison LS, Popescu DL, Anderson ME, et al. Long-term voluntary wheel running does not alter vascular amyloid burden but reduces neuroinflammation in the Tg-SwDI mouse model of cerebral amyloid angiopathy. Journal of Neuroinflammation. 2019;16(1):1–15. doi:10.1186/s12974-019-1534-0

73. Lu X, Moeini M, Li B, et al. Voluntary Exercise Increases Brain Tissue Oxygenation and Spatially Homogenizes Oxygen Delivery in a Mouse Model of Alzheimer’s Disease. Vol 88. Elsevier Inc.; 2020. doi:10.1016/j.neurobiolaging.2019.11.015

74. Lee JH, Stefan S, Walek K, et al. Investigating the correlation between early vascular alterations and cognitive impairment in Alzheimer’s disease in mice with SD-OCT. Biomed Opt Express. 2023;14(4):1494. doi:10.1364/BOE.481826

75. Madsen LS, Parbo P, Ismail R, et al. Capillary dysfunction correlates with cortical amyloid load in early Alzheimer’s disease. Neurobiology of Aging. 2023;123:1–9. doi:10.1016/j.neurobiolaging.2022.12.006

76. Roy TK, Secomb TW. Functional implications of microvascular heterogeneity for oxygen uptake and utilization. Physiological Reports. 2022;10(10):e15303. doi:10.14814/phy2.15303

77. Pries AR, Secomb TW, Gaehtgens P. Design Principles of Vascular Beds. Circulation Research. 1995;77(5):1017–1023. doi:10.1161/01.RES.77.5.1017

78. Korte N, Barkaway A, Wells J, et al. Inhibiting Ca2+ channels in Alzheimer’s disease model mice relaxes pericytes, improves cerebral blood flow and reduces immune cell stalling and hypoxia. Nat Neurosci. Published online September 18, 2024. doi:10.1038/s41593-024-01753-w

79. Yamaguchi T, Ishikawa T, Imai Y, eds. 2 - Biomechanics of Microcirculation. In: Integrated Nano-Biomechanics. Micro and Nano Technologies. Elsevier; 2018:9–69. 10.1016/B978-0-323-38944-0.00002-4

80. Lu X, Moeini M, Li B, et al. Voluntary exercise increases brain tissue oxygenation and spatially homogenizes oxygen delivery in a mouse model of Alzheimer’s disease. Neurobiology of Aging. 2020;88:11–23. doi:10.1016/j.neurobiolaging.2019.11.015

81. Elia A, Parodi-Rullan R, Vazquez-Torres R, Carey A, Javadov S, Fossati S. Amyloid β induces cardiac dysfunction and neuro-signaling impairment in the heart of an Alzheimer’s disease model. Published online July 11, 2023. doi:10.1101/2023.07.11.548558

82. Wrzesień A, Andrzejewski K, Jampolska M, Kaczyńska K. Respiratory Dysfunction in Alzheimer’s Disease—Consequence or Underlying Cause? Applying Animal Models to the Study of Respiratory Malfunctions. Int J Mol Sci. Published online 2024.

83. Lee JH, Ryan J, Andreescu C, Aizenstein H, Lim HK. Brainstem morphological changes in Alzheimer’s disease. Published online 2015.

84. Dutschmann M, Menuet C, Stettner GM, et al. Upper Airway Dysfunction of Tau-P301L Mice Correlates with Tauopathy in Midbrain and Ponto-Medullary Brainstem Nuclei.

85. Li Z, Peng M, Chen P, et al. Imatinib and methazolamide ameliorate COVID-19-induced metabolic complications via elevating ACE2 enzymatic activity and inhibiting viral entry. Cell Metabolism. 2022;34(3):424–440.e7. doi:10.1016/j.cmet.2022.01.008

86. Shah GN, Price TO, Banks WA, et al. Pharmacological Inhibition of Mitochondrial Carbonic Anhydrases Protects Mouse Cerebral Pericytes from High Glucose-Induced Oxidative Stress and Apoptosis. J Pharmacol Exp Ther. 2013;344(3):637–645. doi:10.1124/jpet.112.201400

87. Price TO, Eranki V, Banks WA, Ercal N, Shah GN. Topiramate treatment protects blood-brain barrier pericytes from hyperglycemia-induced oxidative damage in diabetic mice. Endocrinology. 2012;153(1):362–372. doi:10.1210/en.2011-1638

88. Gonzales JU, Scheuermann BW. Effect of acetazolamide on respiratory muscle fatigue in humans. Respiratory Physiology & Neurobiology. 2013;185(2):386–392. doi:10.1016/j.resp.2012.08.023

89. Meekers E, Dauw J, Martens P, et al. Renal function and decongestion with acetazolamide in acute decompensated heart failure: the ADVOR trial. European Heart Journal. 2023;44(37):3672–3682. doi:10.1093/eurheartj/ehad557

90. Solesio ME, Peixoto PM, Debure L, et al. Carbonic anhydrase inhibition selectively prevents amyloid β neurovascular mitochondrial toxicity. Aging Cell. 2018;17(4). doi:10.1111/acel.12787

91. Fossati S, Giannoni P, Solesio ME, et al. The carbonic anhydrase inhibitor methazolamide prevents amyloid beta-induced mitochondrial dysfunction and caspase activation protecting neuronal and glial cells in vitro and in the mouse brain. Neurobiology of Disease. Published online 2016. doi:10.1016/j.nbd.2015.11.006

92. Eskildsen SF, Gyldensted L, Nagenthiraja K, et al. Increased cortical capillary transit time heterogeneity in Alzheimer’s disease: a DSC-MRI perfusion study. Neurobiology of Aging. 2017;50:107–118. doi:10.1016/j.neurobiolaging.2016.11.004

