## Supplementary for "Carbonic anhydrase inhibitors prevent presymptomatic capillary flow disturbances in a model of cerebral amyloidosis" (from another version of this preprint: version 1, 2024-08-22)

### Supplementary information

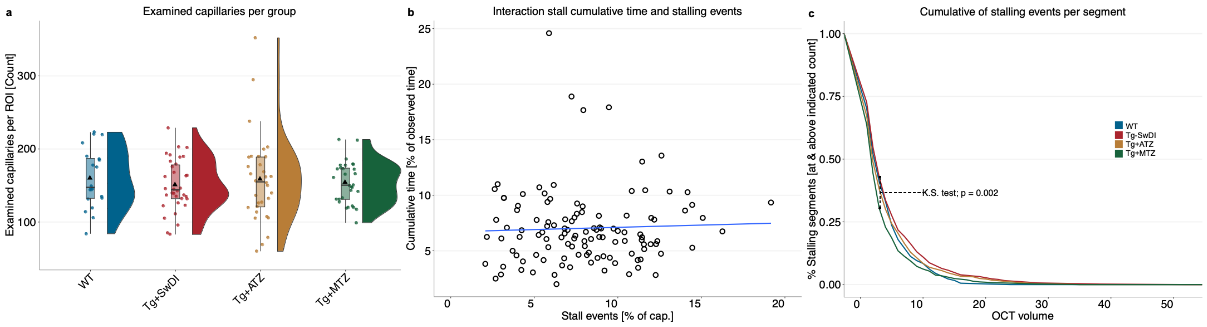

**Supplementary figure 1 | a**, Number of capillaries examined per ROI in each group. **b,** Stall incidence is not correlated with average cumulative stall duration per scanned region (LMM, p = 1). **c**, Average capillary stalling episodes per OCT-A volume (~8.5 seconds per volume). **h**, no difference was observed between groups on the examined capillaries per OCT angiogram. **i**, probability distribution of the fraction of stalled segments and the cumulative duration. Tg+MTZ mice showed fewer capillary segments with large cumulative stalling duration (Kolmogorov-Smirnov test, p = 0.002). Error bar = 100 µm. p-values: * p ≤ 0.05, ** p ≤ 0.01.

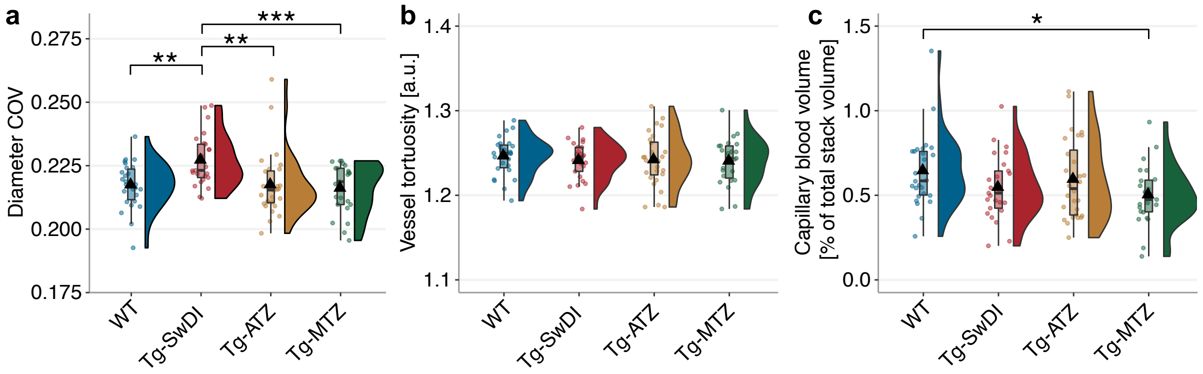

**Supplementary figure 2** | **a,** Diameter coefficient of variance (COV) was estimated as the ratio of STD and diameter. Tg-SwDI vs. WT, LMM, diff = 0.01 ± 0.003, t(51)=3.5, p = 0.002; Tg-SwDI vs. Tg+ATZ, LMM, diff. = –0.01 ± 0.003, t(102) = –3.3, p = 0.002; Tg-SwDI vs. Tg+MTZ, diff. = –0.01 ± 0.003, t(102) = –3.5, p < 0.001). **b**, vessel tortuosity was estimated as the Euclidean distance between two segments ends. **c**, capillary blood volume (CapBV) was estimated as the percentage of volume occupied by the capillary segments in the total z-stack. WT vs. Tg+MTZ, LMM, diff. = –0.14 ± 0.07, t(102) = –2.1, p = 0.04).

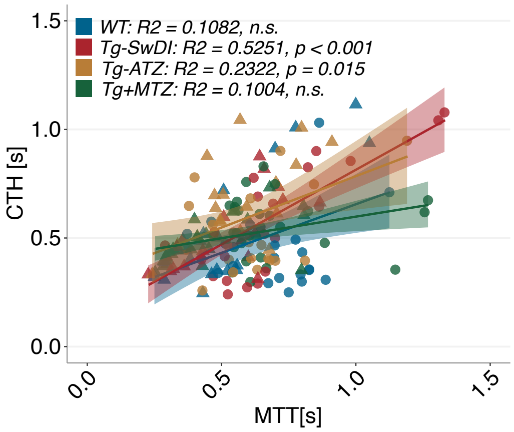

**Supplementary figure 3 |** Regression plot of CTH as function of MTT from the mean estimated among all scanning trials (3 scans per mouse). No linearity is observed in the WT and Tg+MTZ groups, suggesting an active mechanism at the capillary level compensating for a decrease in cerebral blood flow (low MTT).

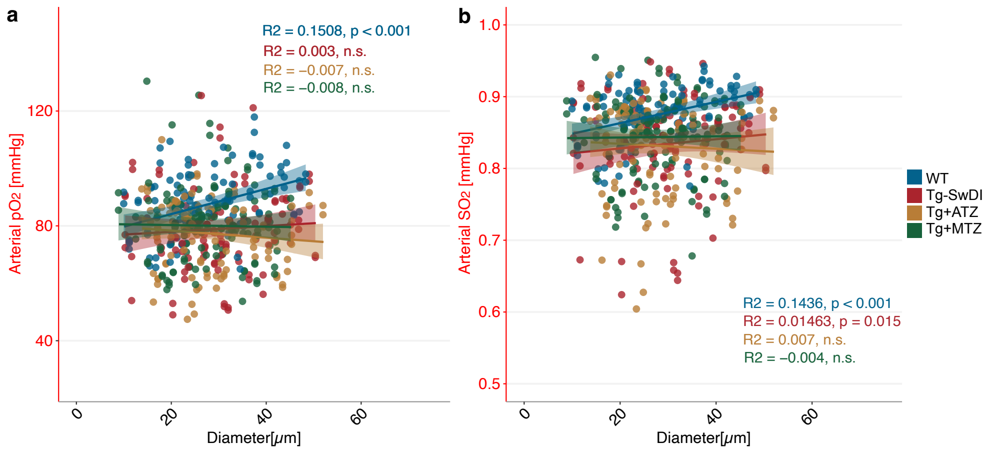

**Supplementary figure 4** | Arterial PO2 **(a)** and SO2 **(b)** as a function of vessel diameter. Linear regression was used to test if diameter predicted PO2 and SO2 in the arterial network in all groups. We found that vessel diameter in the arterial network predicted PO2 ($F(1,222) = 40.6$, $p < 0.001$) and SO2 ($F(1,222) = 38.4$, $p < 0.001$) in the WT group. This prediction was not present in the Tg groups., except for the Tg-SwDI mice where diameter weakly predicted SO2 ($F(1,205) = 4.058$, $p = 0.046$).

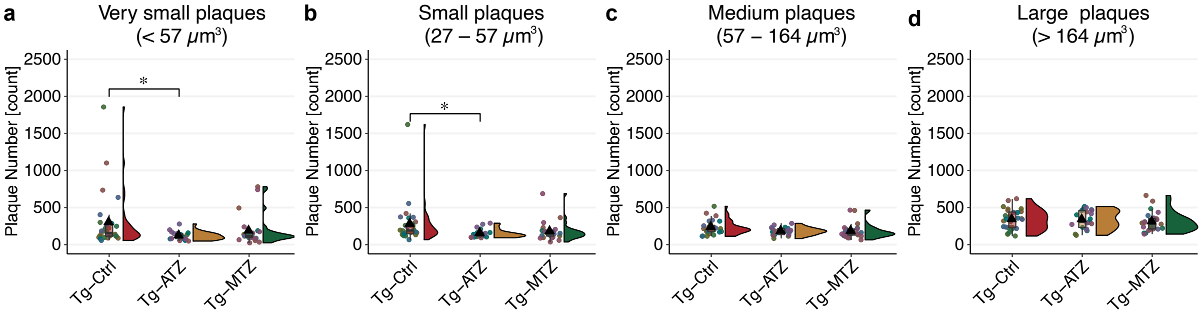

**Supplementary figure 5** | Subpopulation of plaques were done based on quantiles and classified as very small plaques (**a**; LMM, diff. = –178 ± 87 plaques/mm3, t(73) = –2.0, p = 0.048), small plaques (**b**; LMM, diff. = –123 ± 59 plaques/mm3, t(73) = –2.1, p = 0.044), medium size plaques (**c**), and large plaques (**d**).

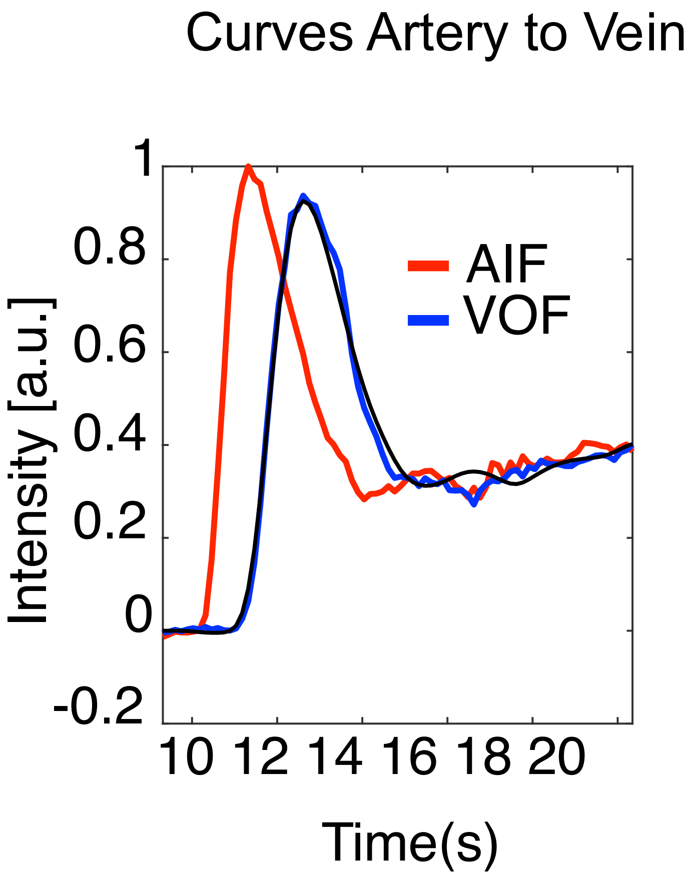

**Supplementary figure 6 |** Visualization of MTT estimated by comparing AIF and VOF after time shifting the curves by the fitted MTT value (delay).

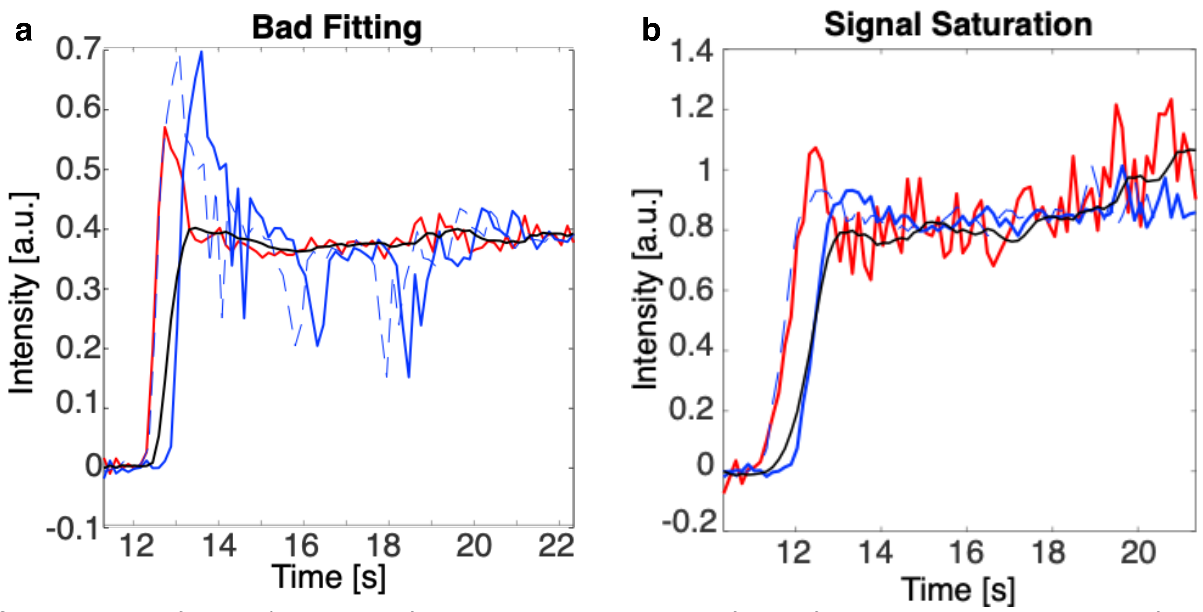

**Supplementary figure 7** | Examples of rejected curves. **a.**, Example for bad fitting. Estimated venous output function (VOF) does not fit with recorded VOF. **b.**, Example for signal saturation.

**Supplementary table 1.** Capillary stalling parameters across experimental groups

| | WT *(n = 20 ROIs in 11 mice)* | Tg-SwDI *(n = 34 ROIs in 18 mice)* | Tg+ATZ *(n = 32 ROIs in 18 mice)* | Tg+MTZ *(n = 30 ROIs in 15 mice)* |
|---|---|---|---|---|
| Stalling events (% of all capillaries) | 5.89 ± 3.37% | 8.4 ± 3.4% | 9.1 ± 3.3% | 7.9 ± 3.4% |
| Stalling events per OCT-A volume | 0.4 ± 0.2% | 0.7 ± 0.4% | 0.6 ± 0.4% | 0.5 ± 0.3% |
| Cumulative time | 6.9 ± 2.5% | 8.4 ± 4.3% | 6.8 ± 3.6% | 5.9 ± 1.9% |
| Examined capillary segments | 160 ± 41 | 150 ± 35 | 158 ± 60 | 154 ± 29 |

**Supplementary table 2.** Capillary hemodynamics from single capillary line scans.

| | WT *(n = 6 mice)* | Tg-SwDI *(n = 7 mice)* | Tg+ATZ *(n = 10 mice)* | Tg+MTZ *(n =9 mice)* |
|---|---|---|---|---|
| Capillaries segments | 113 | 120 | 201 | 149 |
| Capillary diameter (μm) | 3.31 ± 0.45 | 3.21 ± 0.55 | 3.38 ± 0.52 | 3.65 ± 0.52 |
| RBCv (mm/s) | 0.98 ± 0.46 | 0.74 ± 0.36 | 1.00 ± 0.55 | 1.09 ± 0.58 |
| RBCv COV (RBCv SD / RBCv mean) | 0.13 ± 0.12 | 0.12 ± 0.12 | 0.10 ± 0.07 | 0.10 ± 0.05 |
| Cell flux (cell/s) | 47 ± 17 | 42 ± 15 | 51 ± 21 | 56 ± 24 |
| Flux COV (Flux SD / Flux mean) | 0.14 ± 0.07 | 0.16 ± 0.16 | 0.13 ± 0.07 | 0.15 ± 0.07 |
| LD (cells/mm) | 54 ± 18 | 63 ± 22 | 26 ± 19 | 58 ± 21 |
| LD COV (LD SD / LD mean) | 0.14 ± 0.09 | 0.14 ± 0.10 | 0.13 ± 0.09 | 0.13 ± 0.08 |

**Supplementary table 3.** Capillary morphometrics from analysis of vascular stacks.

| | WT *(n = 26 ROIs in 13 mice)* | Tg-SwDI *(n = 25 ROIs in 13 mice)* | Tg+ATZ *(n = 28 ROIs in 14 mice)* | Tg+MTZ *(n = 23 ROIs in 12 mice)* |
|---|---|---|---|---|
| Capillaries segments | 4778 | 4566 | 4752 | 3676 |
| Mean Diameter per ROI (μm) | 5.69 ± 1.49 | 5.74 ± 1.51 | 5.73 ± 1.55 | 5.85 ± 1.60 |
| Mean STD per ROI (μm) | 1.17 ± 0.06 | 1.22 ± 0.08 | 1.17 ± 0.08 | 1.17 ± 0.05 |
| Mean CV per ROI | 0.22 ± 0.01 | 0.23 ± 0.01 | 0.22 ± 0.01 | 0.21 ± 0.01 |
| Capillary Tortuosity | 1.25 ± 0.02 | 1.24 ± 0.02 | 1.24 ± 0.03 | 1.23 ± 0.03 |
| Capillary Blood Volume (% of total z-stack volume) | 0.64 ± 0.22 | 0.54 ± 0.19 | 0.59 ± 0.24 | 0.50 ± 0.18 |
| Capillary Density (μm/μl) | 204911 ± 70264 | 171590 ± 59126 | 192583 ± 68503 | 227284 ± 72370 |

**Supplementary table 4.** MTT and CTH group values

| | MTT | *CTH* |
|---|---|---|
| S1: Artery to Vein | | |
| WT (n = 20) | 0.71 ± 0.18 | 0.44 ± 0.18 |
| Tg-SwDI (n = 21) | 0.70 ± 0.26 | 0.57 ± 0.26 |
| Tg+ATZ (n = 20) | 0.62 ± 0.19 | 0.52 ±0.19 |
| Tg+MTZ (n = 29) | 0.70 ± 0.22 | 0.54 ± 0.14 |
| S2: Arteriole to Venule | | |
| WT (n = 20) | 0.52 ± 0.22 | 0.52 ± 0.26 |
| Tg-SwDI(n = 18) | 0.50 ± 0.19 | 0.51 ± 0.17 |
| Tg+ATZ (n = 18) | 0.56 ± 0.17 | 0.67 ± 0.20 |
| Tg+MTZ (n = 26) | 0.48 ± 0.18 | 0.48 ± 0.11 |

**Supplementary table 5.** Pial vessel diameter in µm

| | WT<br>*(n = )* | Tg-SwDI<br>*(n =)* | Tg+ATZ<br>*(n =)* | Tg+MTZ<br>*(n =)* |
|---|---|---|---|---|
| Arteries | 27.78 ± 9.95<br>*(n = 52)* | 30.01 ± 10.72<br>*(n = 54)* | 27.84 ± 9.33<br>*(n = 57)* | 28.98 ± 9.35<br>*(n = 63)* |
| Diving arterioles | 17.97 ± 5.18<br>*(n = 51)* | 17.54 ± 6.15<br>*(n = 54)* | 16.28 ± 3.37<br>*(n = 67)* | 19.72 ± 8.8<br>*(n = 66)* |
| Upstream venules | 19.14 ± 7.10<br>*(n = 96)* | 18.41 ± 6.97<br>*(n = 89)* | 16.64 ± 5.89<br>*(n = 87)* | 18.90 ± 6.50<br>*(n = 115)* |
| Veins | 38.06 ± 18.72<br>*(n = 56)* | 37.92 ± 18.72<br>*(n = 53)* | 41.10 ± 23.97<br>*(n = 44)* | 37.46 ± 19.64<br>*(n = 64)* |

**Supplementary table 6.** Mean PO2 (mmHg), SO2 (fractional) and OEF (fractional).

| | WT *(n = )* | Tg-SwDI *(n =)* | Tg+ATZ *(n =)* | Tg+MTZ *(n =)* |
|---|---|---|---|---|
| **PO2** | | | | |
| Arteries | 87.44 ± 11.2 *(n = 111)* | 79.0 ± 14.7 *(n = 101)* | 77.4 ± 11.9 *(n = 114)* | 80.0 ± 13.2 *(n = 129)* |
| Diving arterioles | 82.5 ± 13.4 *(n = 113)* | 80.8 ± 19.2 *(n = 106)* | 74.5 ± 11.3 *(n = 137)* | 74.1 ± 15.7 *(n = 145)* |
| Ascending venules | 47.9 ± 9.1 *(n = 172)* | 44.5 ± 10.3 *(n = 149)* | 43.7 ± 8.7 *(n = 163)* | 49.2 ± 8.6 *(n = 210)* |
| Veins | 48.3 ± 8.1 *(n = 99)* | 48.1 ± 9.6 *(n = 97)* | 43.7 ± 9.7 *(n = 88)* | 49.1 ± 7.3 *(n = 186)* |
| **SO2** | | | | |
| Arteries | 0.87 ± 0.04 *(n = 111)* | 0.83 ± 0.07 *(n = 101)* | 0.83 ± 0.06 *(n = 114)* | 0.84 ± 0.05 *(n = 129)* |
| Diving arterioles | 0.85 ± 0.06 *(n = 113)* | 0.83 ± 0.09 *(n = 106)* | 0.82 ± 0.06 *(n = 137)* | 0.81 ± 0.08 *(n = 145)* |
| Ascending venules | 0.59 ± 0.12 *(n = 172)* | 0.54 ± 0.15 *(n = 149)* | 0.54 ± 0.13 *(n = 169)* | 0.61 ± 0.11 *(n = 210)* |
| Veins | 0.60 ± 0.10 *(n = 99)* | 0.60 ± 0.12 *(n = 97)* | 0.54 ± 0.15 *(n = 88)* | 0.62 ± 0.09 *(n = 126)* |
| **OEF** | | | | |
| Arteries to veins | 0.30 ± 0.11 *(n = 26)* | 0.31 ± 0.11 *(n = 25)* | 0.36 ± 0.15 *(n = 28)* | 0.28 ± 0.07 *(n = 34)* |
| Diving arterioles to ascending venules | 0.29 ± 0.11 *(n = 26)* | 0.33 ± 0.12 *(n = 25)* | 0.35 ± 0.14 *(n = 28)* | 0.24 ± 0.09 *(n = 34)* |

**Supplementary table 7. ELISA for amyloid load**

| | WT<br>*(n = )* | Tg-SwDI<br>*(n =)* | Tg+ATZ<br>*(n =)* | Tg+MTZ<br>*(n =)* |
|---|---|---|---|---|
| Aβ-40 (pg/ml) - Cortex | 63 ± 61 (n = 12) | 85867 ± 19852 (n = 9) | 68450 ± 19852 (n = 9) | 73858 ± 20333 (n = 9) |
| Aβ-40 (pg/ml) - Hippocampus | 126 ± 161 (n = 9) | 75875 ± 29062 (n = 9) | 51824 ± 14892 (n = 9) | 73377 ± 26156 (n = 9) |
| Aβ-42 (pg/ml) – Cortex | 260 ± 242 (n = 12) | 1883 ± 668 (n = 9) | 1320 ± 280 (n = 9) | 1426 ± 233 (n = 9) |
| Aβ-42 (pg/ml) - Hippocampus | 338 ± 329 (n = 9) | 2289 ± 1158 (n = 9) | 1428 ± 230 (n = 9) | 1723 ± 364 (n = 9) |
